# A three amino acid sequence gates protein stability control of bHLH104 and iron uptake in *Arabidopsis thaliana*

**DOI:** 10.1101/2025.11.18.689088

**Authors:** Dibin Baby, Sarah Weldi, Marie Knopf, Vinitha Venkadasamy, Jessica Lee Erickson, Petra Bauer

**Author notes:** **Corresponding author:** Petra Bauer.

## Abstract

Iron (Fe) homeostasis is regulated to prevent iron imbalance, with the help of redundant basic helix-loop-helix (bHLH) IVc transcription factors that can be controlled by Fe-binding E3 ligases such as BRUTUS (BTS) in *Arabidopsis thaliana*. However, knowledge gaps remain to fully explain the mechanistic basis of this redundant protein interaction module. The C-terminus of bHLH104 interacts with BTS. Structural predictions suggest involvement of the three terminal amino acids, proline-alanine-alanine (PAA). However, the importance of the PAA short sequence for post-translational regulation of bHLH104 and resulting plant phenotypes has not been experimentally tested. Here, we demonstrate that transgenic plants expressing a bHLH104 variant lacking PAA (*b104^dPAA*) constitutively upregulated Fe acquisition and Fe transport genes and consequently accumulated Fe in contrast to wild-type bHLH104 protein-expressing plants. The extent of Fe accumulation in independent b104^dPAA lines correlated positively with b104^dPAA protein abundance. In contrast to the wild-type bHLH104 form, there was no indication that b104^dPAA interacted with BTS or was ubiquitinated by it. Together, these findings corroborated that the PAA sequence is required for post-translational control of bHLH104 by BTS. Hence, targeted manipulation of bHLHIVc protein PAA sequence may represent a strategy for crop biofortification.

**Plain language summary:** Plants tightly control iron levels. In *Arabidopsis thaliana*, transcription factor bHLH104 steers iron uptake. E3 ligases like BTS regulate bHLH104. We show that a three–amino acid terminal sequence of bHLH104 is needed for its targeting by the BTS pathway. bHLH104 lacking PAA leads to iron accumulation. Targeting this short sequence can be a strategy for crop biofortification.

## Introduction

Iron (Fe) is an essential micronutrient required for plant growth and development. It plays critical roles in photosynthesis, respiration, chlorophyll biosynthesis, and redox reactions (Briat et al., 2007; Jeong and Guerinot, 2009). Although Fe is abundant in the Earth’s crust, it is mostly present in the ferric (Fe³⁺) form, which is poorly soluble and difficult for plant roots to absorb—especially in alkaline or calcareous soils (Marschner, 2012). Excess Fe is toxic since it generates reactive oxygen species (ROS) (Panda and Yamamoto, 2019). Therefore, plants must tightly regulate Fe uptake, distribution, and storage to maintain internal balance.

In *Arabidopsis thaliana*, a group of basic helix–loop–helix (bHLH) IVc transcription factors—including bHLH34, bHLH104, ILR3 (bHLH105), and bHLH115—acts as an early regulatory module in the Fe deficiency (-Fe) response. These proteins activate the expression of internal Fe allocation and Fe deficiency response regulatory genes in roots and shoots (Schwarz and Bauer, 2020). Among their targets are also genes coding for bHLH Ib transcription factors that, together with the bHLH transcription factor FIT (FER-LIKE IRON DEFICIENCY-INDUCED TRANSCRIPTION FACTOR), induce root Fe uptake genes such as *IRT1* and *FRO2* (Long et al., 2010; Selote et al., 2015; Zhang et al., 2015; Li et al., 2016; Liang et al., 2017).

The regulation of the redundant bHLH IVc transcription factors is not fully understood. Notably, transcript levels of *BHLH IVc* genes do not change substantially under -Fe conditions (Zhang et al., 2015; Li et al., 2016; Liang et al., 2017), indicating that they are activated during the -Fe response primarily at protein level. It is proposed that bHLH IVc proteins are subject to post-translational regulation via the 26S proteasome, mediated by the E3 ubiquitin ligases, BRUTUS (BTS)/BTS-LIKE (BTSL) proteins (Zhao et al., 2026). However, this has not yet been demonstrated experimentally for all bHLH IVc proteins. It was shown that bHLH105/ILR3 and bHLH115 are recognized, ubiquitinated and degraded by BTS and BTSL proteins (Selote et al., 2015; Hindt et al., 2017; Zhao et al., 2026). Control by E3 ligases of the BTS/BTSL-type is particularly intriguing since stability of the BTS/BTSL E3 ligases is controlled by their N-terminus which contains hemerythrin domains for Fe and oxygen sensing (Selote et al., 2015; Pullin et al., 2025). Additionally, Fe and Zn binding to C-terminal zinc finger domains (CHY, CTCHY, RING) and rubredoxin-type folds of the BTS rice ortholog HEMERYTHRIN MOTIF-CONTAINING REALLY INTERESTING NEW GENE AND ZINC-FINGER PROTEIN (HRZ) may regulate its protein stability and activity, suggesting an additional mechanism for metal ion sensing (Shinkawa et al., 2025). Together, these findings support a model in which, under Fe deficiency, altered metal occupancy and redox state of BTS/BTSL domains attenuate E3 ligase activity toward bHLH IVc transcription factors, allowing accumulation of the transcription factors and subsequent activation of Fe uptake genes. This model is further supported by genetic studies demonstrating that loss of BTS (Selote et al., 2015; Long et al., 2010) or BTSL1/BTSL2 (Hindt et al., 2017; Rodríguez-Celma et al., 2019) results in activation of Fe deficiency responses. However, due to technical limitations and the redundancy among the involved proteins, there remain knowledge gaps.

One bHLH IVc group protein is bHLH104 (Zhang et al., 2015; Li et al., 2016). Phylogenetic analysis places bHLH104 together with bHLH34 in a distinct clade from bHLH105 and bHLH115, and individual bHLH IVc members show distinct expression patterns (Gao et al., 2019; Gao et al., 2024; Jiang et al., 2024), indicating that regulatory mechanisms cannot be directly extrapolated between members. T-DNA loss-of-function lines of *bHLH104* exhibit an Fe deficiency-sensitive phenotype (Li et al., 2016), which is expected for an upstream regulator of the Fe deficiency response. The protein-level regulatory mechanisms controlling bHLH104 function remain incompletely characterized. The C-terminus of bHLH104 can interact with BTS/L proteins (Lichtblau et al., 2022). It possesses the sequence PAA at the very C-terminal end. In our previous study, we showed that deleting a 25-amino-acid-long stretch at the C terminus abolished bHLH104 interaction with BTS (Lichtblau et al., 2022). Other studies supported the implication of this C-terminal region in regulation of bHLH IVc proteins. (Li et al., 2021) demonstrated that substitution of the terminal alanine to valine in bHLH105 and bHLH115 abolished interaction with BTS in yeast two-hybrid assays and increased protein stability in transient expression assays. (Sharma and Yeh, 2020) identified an ethyl methanesulfonate (EMS)-induced C-terminal alanine-to-valine substitution in bHLH34 that conferred increased protein stability and metal accumulation in plants, but a connection to BTS was not established. A C-terminal sequence with terminal PAA is conserved in IRON MAN (IMA)/FE-UPTAKE-INDUCING PEPTIDE (FEP) signaling peptides. Interestingly, (Lichtblau et al., 2022) found that the C-terminal portion of IMA1 small proteins is needed for IMA1 to interact with the BTSL1 C-terminus. Moreover, they described that intact IMA1, but not IMA1 with a C-terminal deletion, disturbed interaction between C-terminal protein fragments of BTSL1 and either bHLH34, bHLH104, or bHLH115, but not ILR3. (Li et al., 2021) showed that substitution of the terminal alanine to valine in IMA3 abolished BTS interaction, further demonstrating the importance of this terminal residue for protein recognition by BTS. Structural predictions suggested that the conserved three–amino acid short sequence—proline–alanine–alanine (PAA)—of a bHLH transcription factor may bind at the BTS-binding interface (Lichtblau et al., 2022). In all, the BTS and BTSL recognition may thus be narrowed down to PAA/PVA. Hereafter, we refer to this as the “PAA short sequence”.

Although BTS-mediated ubiquitination has been demonstrated for *A. thaliana* bHLH105 and bHLH115 (Li et al., 2021; Zhao et al., 2026), direct ubiquitination of bHLH104 by BTS has not been shown, nor has the requirement of PAA for this interaction been established. Genetic evidence supports that bHLH104 acts downstream of BTS, as loss of bHLH104 partially suppresses the Fe deficiency tolerance of the *bts-2* mutant (Zhang et al., 2015). Yet, the biochemical basis of this regulation remained unresolved.

The key open questions and knowledge gaps of this study were to examine whether targeting the bHLH104 PAA short sequence in a deletion study would cause any Fe- related plant growth phenotypes. Further, if so, we intended to study whether the underlying mechanism could be explained by disturbed protein abundance. Moreover, we aimed to examine the relevance of PAA for protein interaction with BTS and BTS-mediated ubiquitination. Our work provides evidence that the PAA short sequence is indeed critical for regulating bHLH104 stability and Fe homeostasis across plant development, and our data highlight a potential new strategy for Fe biofortification.

## Results

### Plants with bHLH104 lacking PAA (*b104^dPAA*) have distinct growth phenotypes

We hypothesized that the PAA short sequence plays a role in the functionality of bHLH104 in plants (Figure 1). To test this, we studied transgenic *Arabidopsis thaliana* lines in the wild-type Col-0 background expressing either regular full-length mTurquoise2 fluorophore-tagged bHLH104 (hereafter *pb104::b104*) or a mutant deletion variant lacking the conserved PAA short sequence (hereafter *pb104::b104^dPAA*), both driven by the native *BHLH104* promoter (Figure 2A). The N-terminal mTurquoise2-tagged *-b104^dPAA* complemented the *bhlh104-1* seedling -Fe sensitivity phenotype indicating functionality of the fusion protein (Supplementary Figure S1).

**Figure 1.**
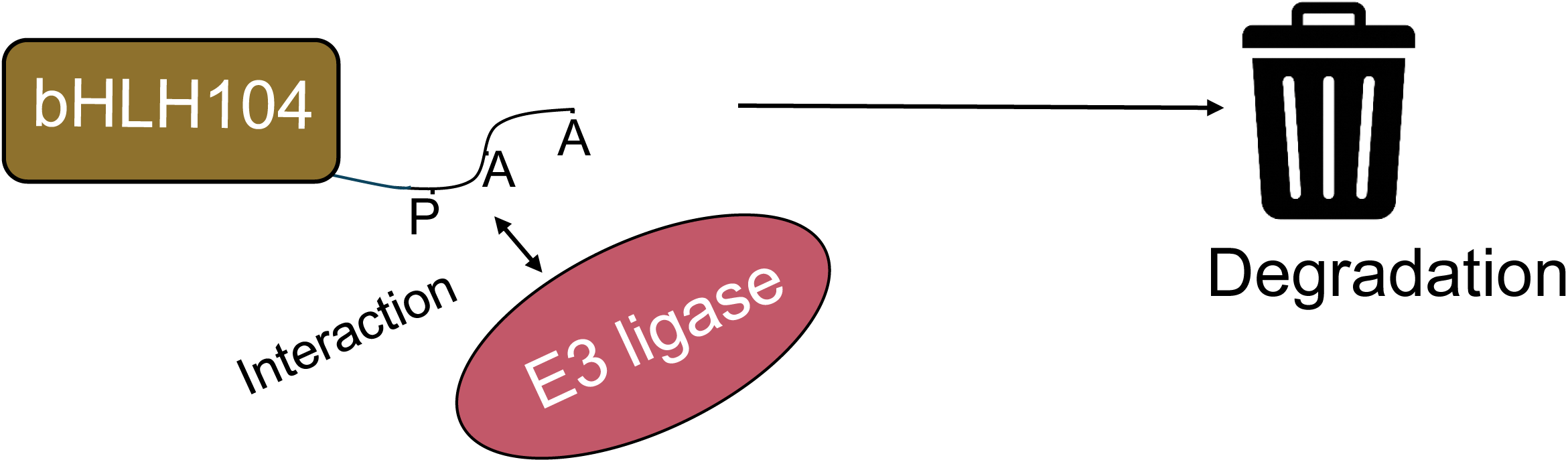
Conceptual model of bHLH104 regulation by its conserved PAA sequence. In the full-length bHLH104 protein, the PAA sequence forms the interaction interface with an E3 ligase, likely enabling ubiquitination and proteasomal degradation of bHLH104.

**Figure 2.**
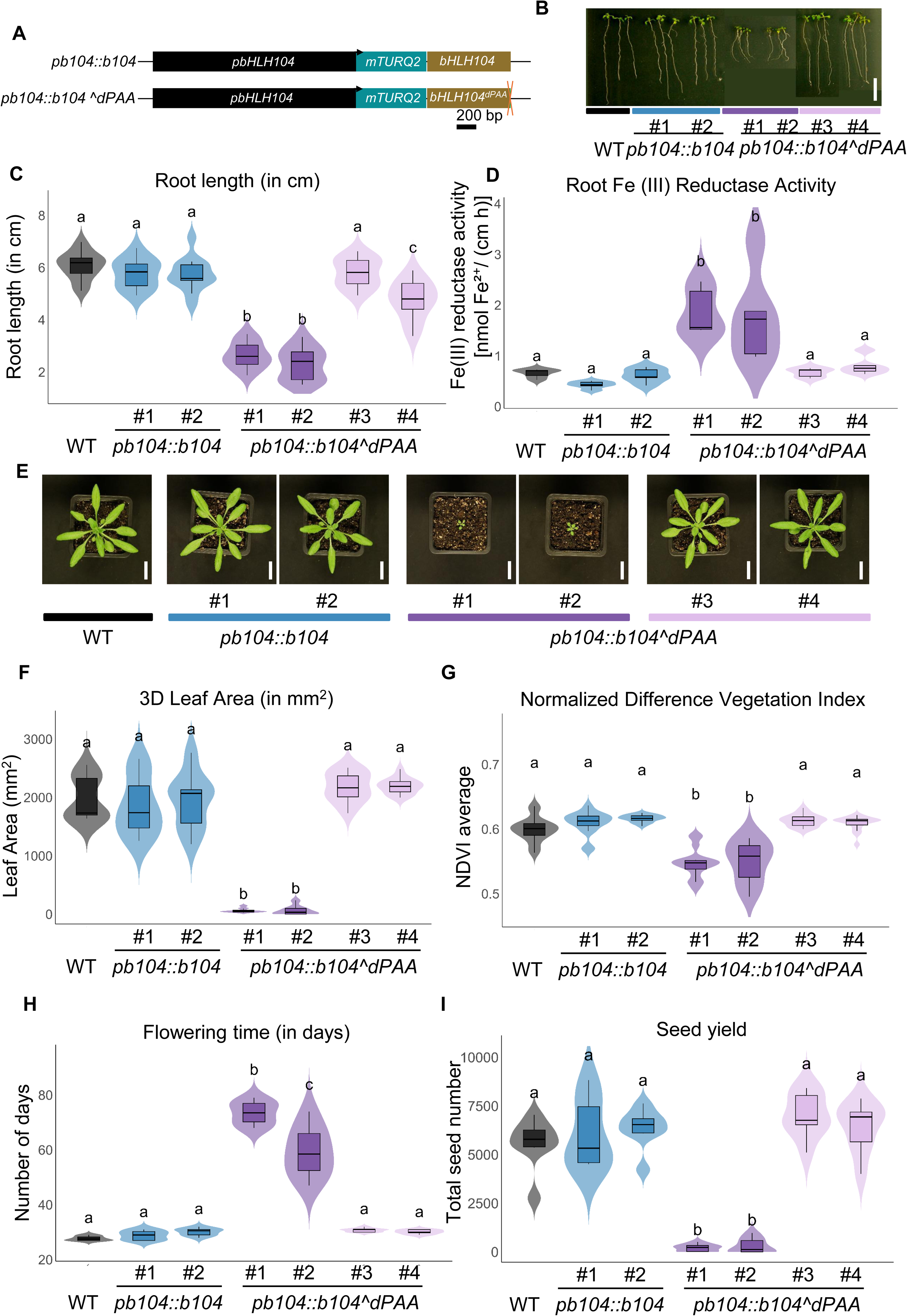
Transgenic expression of bHLH104 lacking the PAA sequence leads to growth alterations and physiological changes in Arabidopsis. (A) Schematic representation of constructs used to generate transgenic lines expressing either full-length bHLH104 (pb104::b104) or a C-terminal deletion variant lacking the conserved three–amino acid PAA sequence (pb104::b104^dPAA). Both constructs are driven by the native bHLH104 promoter (2,069 bp) and fused at the N-terminus to an mTurquoise2 fluorescent tag. The deleted PAA sequence in b104^dPAA is indicated by a red cross at the C-terminus. Scale bar = 200 bp. (B) Representative images of 10-day-old seedlings grown on +Fe medium (scale bar = 2 cm). (C) Primary root length of 12-day-old seedlings grown vertically on Hoagland medium supplemented with Fe. (D) Fe reductase activity in roots of 10-day-old seedlings grown under +Fe. Activity was quantified spectrophotometrically as Fe(II)–ferrozine complex formation and normalized to total root length. (E) Representative images of four-week-old soil-grown plants (scale bar = 2 cm). Shown are Col-0 (WT), two independent pb104::b104 lines (#1, #2), and four independent pb104::b104^dPAA lines (#1–4). (F–G) Quantitative analysis of aboveground phenotypes using the Phenospex PlantEye system: (F) 3D leaf area (mm²); (G) NDVI (Normalized Difference Vegetation Index) (H) Flowering time, recorded as days to bolting. (I) Seed number per plant, estimated from total seed weight based on mean weight per 100 seeds. Data represent means ± SD (n = 7–12 biological replicates per line). Statistical analysis was performed using two-way ANOVA followed by Tukey’s post hoc test. Different letters indicate statistically significant differences at p < 0.05.

Interestingly, phenotypic variation was visible in the T1 population of the *pb104::b104^dPAA* transgenics, indicating a dominant transgene effect: While the majority of plants displayed severely reduced growth with brown necrotic spots on leaves, some plants appeared indistinguishable from wild type (WT) (Supplementary Figure S2). To capture these contrasting phenotypes, we selected and propagated four *pb104::b104^dPAA* lines, two with severe phenotypes (#1, #2) and two with no obvious macroscopic phenotype (#3, #4), in addition to *pb104::b104 #1* and *#2* controls.

To quantify phenotypic variation across the life cycle, detailed morphological and physiological assessments were made with T3 plants selected for transgene presence by genotyping PCR and grown on agar plates or soil. In 12-day-old seedlings of *pb104::b104^dPAA* lines *#1* and *#2*, we found significantly shorter primary roots than in seedlings of #3, #4 or in *pb104::b104* control lines (Figure 2B, 2C). Root Fe(III) reductase activity is necessary for reduction-based Strategy I Fe uptake (Yi and Guerinot, 1996). Ferric reductase activity measurements revealed that *pb104::b104^dPAA* lines *#1* and *#2* had higher Fe reductase activity compared to #3, #4 and the controls (Figure 2D). The machine-aided Phenospex PlantEye phenotyping platform can reliably depict minor variation in leaf color and rosette size in *A. thaliana* (Knopf and Bauer, 2025). By four weeks, *pb104::b104^dPAA* lines *#1* and *#2* displayed smaller rosette area (Figures 2E, 2F) and low normalized difference vegetation index (NDVI) values (Figure 2G), which indicate lower chlorophyll content, lower photosynthetic efficiency, and reduced growth performance compared with other lines. Flowering was delayed compared to the other lines (Figure 2H) and seed yield was significantly lower for *pb104::b104^dPAA* lines *#1* and *#2* compared to the other lines (Figure 2I).

Reduced root and shoot growth, as observed in plants of *pb104::b104^dPAA* line #2, can be a consequence of Fe toxicity (Naranjo-Arcos et al., 2017). Thus, we tested whether the growth defects observed in *pb104::b104^dPAA* #2 plants were alleviated in conditions of Fe deficiency. We grew the representative lines on alkaline calcareous soil (ACS; pH 8.1), which provides Fe-limiting conditions (Supplementary Figure S3; Knopf and Bauer, 2025). Notably, *pb104::b104^dPAA line #2*, which exhibited pronounced growth defects in control soil (pH 5.9), showed increased 3D leaf area in ACS, indicating partial recovery under Fe-limited ACS conditions (Supplementary Figure S3C). NDVI values, which were low in control soil, were restored to near-WT levels in ACS, suggesting improved leaf health (Supplementary Figure S2D). While flowering remained delayed relative to WT, it occurred earlier in ACS than in control soil (Supplementary Figure S3E). These findings show that the growth defects observed in control soil are largely alleviated when Fe availability is limited, linking the b104^dPAA-associated phenotypes to Fe status.

Together, these results show that removal of the PAA short sequence in bHLH104 can lead to drastic dominant growth phenotypes across the life cycle. To explore whether these phenotypes reflect altered Fe homeostasis, we next examined tissue Fe content and expression of Fe-responsive genes in both seedlings and plants at the reproductive stage. For further detailed experiments, we focused on two representative *pb104::b104^dPAA* lines, namely a strong line, #2, and a weak phenotype line, #3.

### b104^dPAA Can Cause Constitutive Iron Uptake and Transcriptional Misregulation of Fe Deficiency Response genes in Seedlings

After a few days of germination, *A. thaliana* seedlings rely on external Fe supply. To assess whether b104^dPAA affects Fe homeostasis during early development, we measured total Fe content via ICP-MS and analyzed gene expression profiles in 10-day-old seedlings of WT, control and contrasting *pb104::b104^dPAA* lines #2 and #3 in both +Fe and -Fe conditions (Figure 3A, B). As an additional control, we used *pUBQ10::b104* lines expressing full-length bHLH104 constitutively from the strong *UBIQUITIN10* promoter, allowing us to assess whether transcript levels alone affect Fe homeostasis. Morphologically, *pUBQ10::b104* lines #1 and #2 are indistinguishable from WT, except for slightly shorter roots (Supplementary Figure S4).

**Figure 3.**
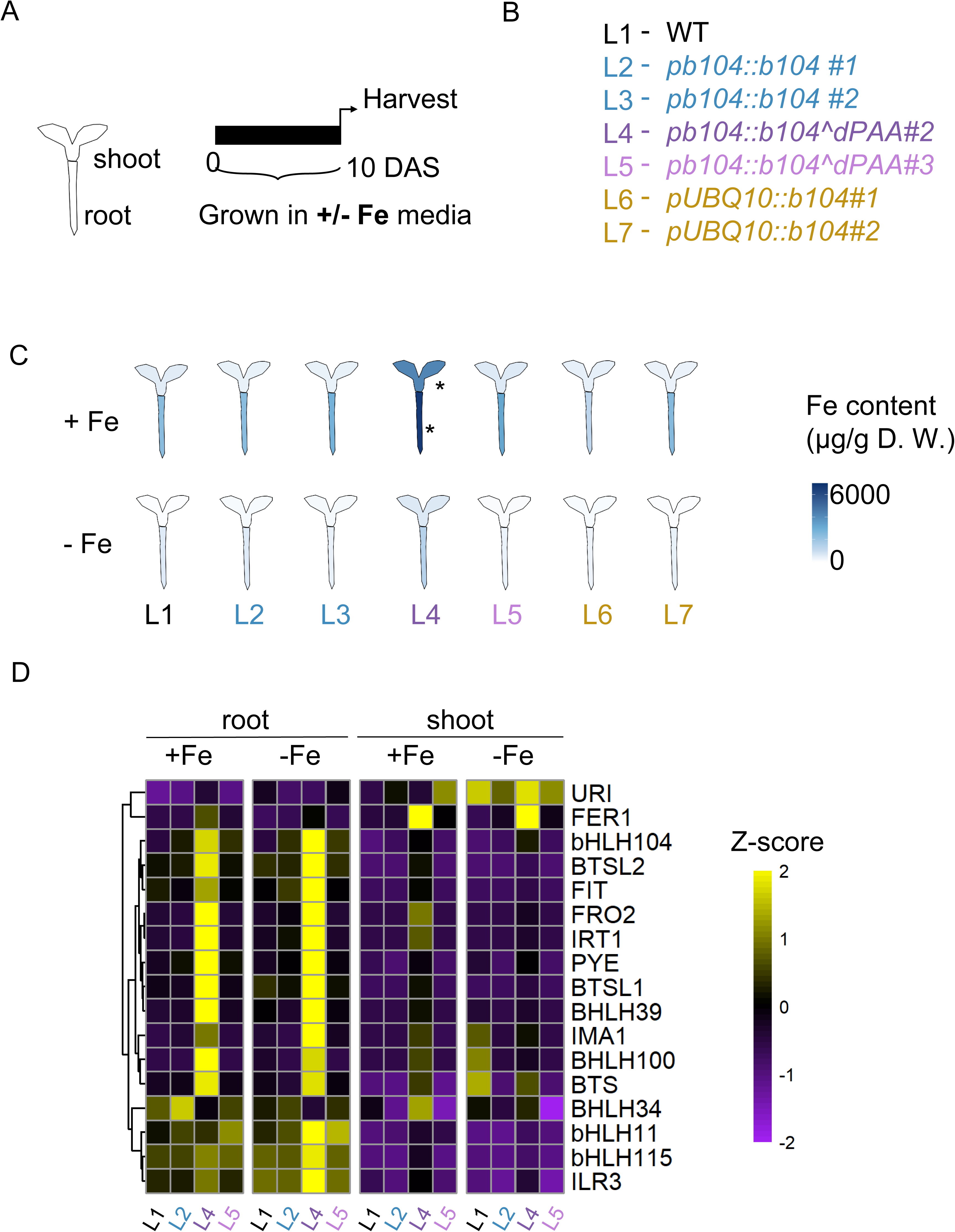
Transgenic lines expressing bHLH104^dPAA exhibit increased iron accumulation and altered iron homeostasis gene expression in seedling stage. (A) Schematic representation of the experimental design. Arabidopsis seedlings were grown for 10 days after sowing (DAS) under either iron-sufficient (+Fe) or iron-deficient (–Fe) conditions. Shoot and root tissues were collected separately for ICP-MS and gene expression analyses. (B) Genotypes used in this study. Abbreviations of lines used: L1, wild-type Col-0; L2, pb104::mTurquoise2-bHLH104 line #1; L3, pb104::mTurquoise2-bHLH104 line #2; L4, pb104::mTurquoise2-b104^dPAA line #2; L5, pb104::mTurquoise2-b104^dPAA line #3; L6, pUBQ10::mTurquoise2-bHLH104 line #1; L7, pUBQ10::mTurquoise2-bHLH104 line #2.(C) Iron content in shoot and root tissues of seedlings grown under +Fe and –Fe conditions, measured by ICP-MS. Data represent the mean of three biological replicates (n = 3). Heat map intensity reflects iron concentration (µg/g dry weight). Color shading was rendered using the R package ggplantmap, with darker blue indicating higher Fe accumulation. Notably, pb104::b104^dPAA lines show increased Fe accumulation in roots and shoots under +Fe conditions. Asterisks indicate genotypes that differ significantly from WT within the same tissue and treatment condition (p < 0.05), as determined by two-way ANOVA and Tukey’s post hoc test (n = 3 for each group). (D) Clustered heatmaps showing the expression profiles (RNA sequencing) of selected Fe homeostasis genes. Heatmaps were generated in RStudio version 4.4.1 using calculated Z-scores of normalized gene expression values across tissues for both +Fe and –Fe conditions. Purple indicates low expression and yellow indicates high expression.

Elemental profiling revealed that *pb104::b104^dPAA* line #2 accumulated substantially more Fe than all other genotypes under both +Fe and –Fe conditions (Figure 3C). This increase was observed in both roots and shoots, with roots showing higher Fe levels than shoots. In contrast, *pb104::b104^dPAA* line #3 and the *pb104::bHLH104* and *pUBQ10::b104* lines had Fe contents similar to WT. All lines exhibited reduced Fe contents under –Fe, confirming the effectiveness of the -Fe treatment (Figure 3C).

Differences were also detected for manganese (Mn), zinc (Zn), and copper (Cu) levels (Supplementary Figure S5). Uptake of these metal ions is closely connected with Fe homeostasis (Connolly et al., 2002). In *pb104::b104^dPAA* line #2, Zn and Cu were elevated in roots under –Fe, and Cu was increased in shoots. Notably, Zn levels were elevated even under +Fe in *pb104::b104^dPAA* roots compared to WT. Roots of *pUBQ10::b104* lines also showed increased Mn under –Fe compared to WT. Metal concentrations in the other lines were similar to WT. Hence, these findings suggest that *pb104::b104^dPAA* line #2 has increased Fe accumulation and affects the uptake of other micronutrients like Zn and Cu. This highlights the importance of proper regulation of bHLH104 function in maintaining metal homeostasis under varying Fe conditions.

To elucidate the transcriptional response, we analyzed gene expression patterns via RNA-seq following -Fe treatment at the seedling stage in both root and shoot tissues. The highest number of differentially expressed genes (DEGs) was observed in shoot tissue of *pb104:b104^dPAA #2* line, with 4,645 DEGs under +Fe conditions and 2,857 DEGs under -Fe conditions after 10 days of treatment, compared to WT. There was also substantial differential expression in roots, with 1,210 DEGs under +Fe conditions and 111 DEGs under -Fe conditions (Supplementary Figure S6A).

Gene Ontology (GO) analysis revealed that the upregulated genes in shoot tissue of the *pb104:b104^dPAA* #2 plants were enriched in terms related to hypoxia and oxidative stress, while genes associated with photosynthesis were downregulated. In *pb104:b104^dPAA* root tissue, upregulated gene categories were additionally enriched for terms related to metal ion transport, while downregulated categories showed enrichment for cell wall modification-related GO terms (Supplementary Figure S6B). Hence, the *pb104:b104^dPAA* #2 plants indeed show symptoms of Fe overload with associated oxidative stress.

Gene expression profiles can inform on misregulation in mutants and transgenic lines (Schwarz and Bauer, 2020).We examined the expression patterns of selected genes involved in Fe uptake and homeostasis regulation to characterize their specific regulatory expression profiles. Interestingly, Fe acquisition and mobilization genes were constitutively upregulated in pb104::b104^dPAA#2 line regardless of Fe availability at the seedling stage (Supplementary Figure S7). *BHLHIVc* genes, *BHLH104, ILR3*, and *BHLH115*, were upregulated in the roots of the *pb104::b104^dPAA line #2* under +Fe compared to other lines (Figure 3D). Their target genes, such as *BHLH39, BHLH100, IMA1, BTS, BTSL1, BTSL2* and *PYE*, were also upregulated, indicating activation of the Fe deficiency response signaling which includes a negative feedback loop via *BTS, BTSL1, BTSL2* and *PYE* genes. Likewise, the further downstream targets in roots, like Fe acquisition genes *IRT1, FRO2, FIT,* were up-regulated (Figure 3D), consistent with the elevated Fe uptake and Fe content. Interestingly, expression of negative regulators of the Fe deficiency response and of Fe sufficiency markers, such as *BHLH11* and *FER1* were also increased, suggesting a proper response to elevated Fe (Figure 3D). In the shoot tissue, the Fe storage related gene *FER1* was strongly upregulated in *pb104::b104^dPAA line #2* compared to controls, regardless of Fe availability, in agreement with the elevated Fe content in this line (Figure 3D). Targeted expression profiling using reverse transcription quantitative PCR (RT-qPCR) in seedling-stage samples confirmed expression patterns to those observed in the transcriptome analysis (Supplementary Figure S8).

To summarize, the misregulation of Fe response genes indicates that bHLH104 with deleted PAA disrupts normal Fe homeostasis by constitutively activating Fe acquisition while preventing proper feedback inhibition of Fe uptake during seedling development. This results in excessive Fe acquisition and resulting toxicity.

### b104^dPAA Causes Constitutive Fe Uptake and Transcriptional Misregulation at a Reproductive Growth Stage

Plants redistribute nutrients during reproduction to support flowering and seed development (Waters and Grusak, 2008; Pottier et al., 2018). To assess whether b104^dPAA affects Fe homeostasis during later developmental stages, we analyzed 30-day-old flowering Arabidopsis plants grown hydroponically, subjected at 27 days to a 3-day ±Fe treatment (Figure 4A). However, the strong-phenotype plants of *pb104::b104^dPAA #2* were severely growth-compromised in hydroponic culture and had to be excluded from this analysis. Instead, we focused on the weak-phenotype line *pb104::b104^dPAA #3*. Old leaves, young leaves, and roots were collected for gene expression analysis, as described in (Ngigi et al., 2025) (Figure 4C-D).

**Figure 4.**
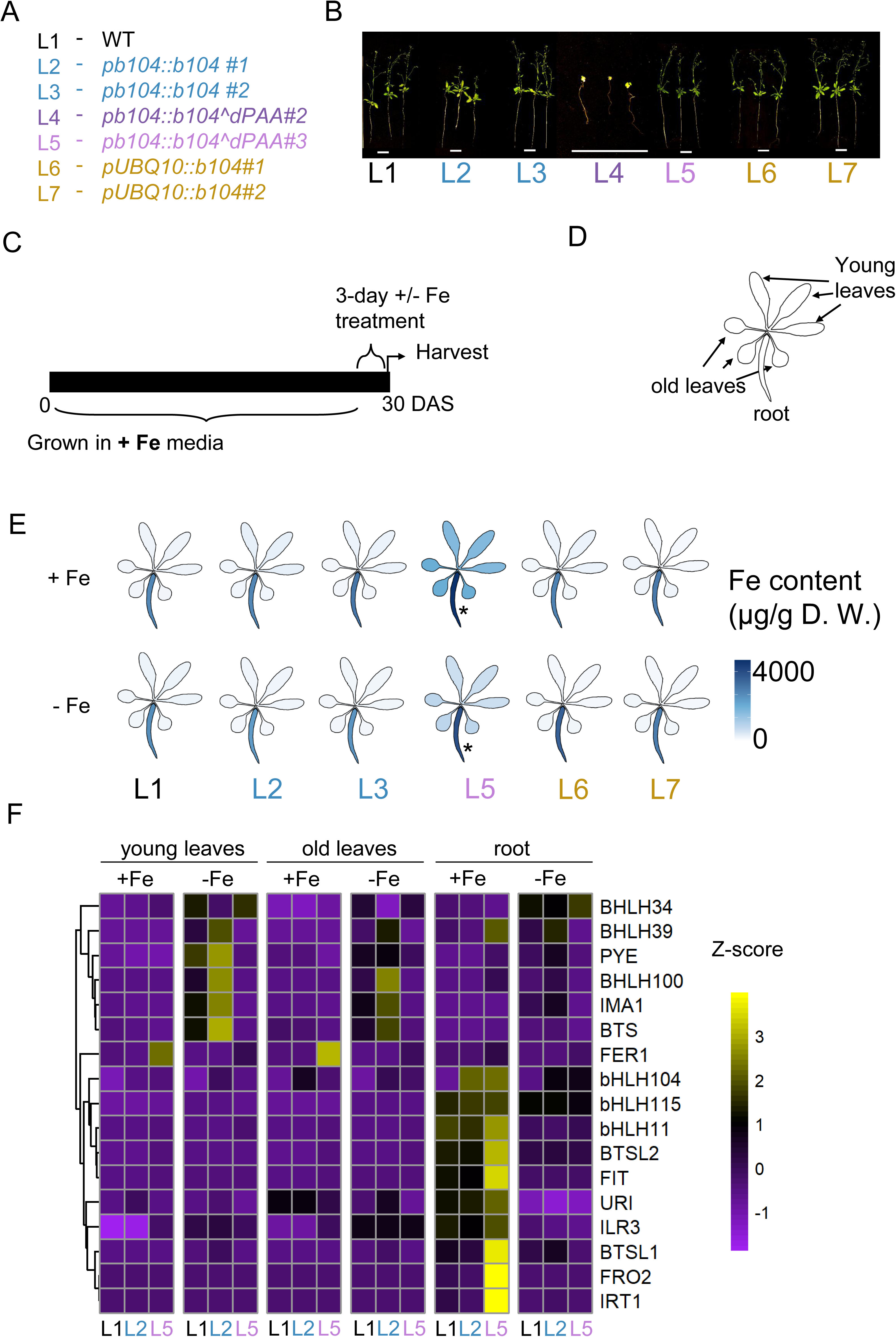
Elevated shoot iron accumulation and altered Fe deficiency responses in reproductive-stage bHLH104^dPAA transgenic lines. (A) Plant lines used In this experiment. Abbreviations of lines used: L1, wild-type Col-0; L2, pb104::mTurquoise2-bHLH104 line #1; L3, pb104::mTurquoise2-bHLH104 line #2; L4, pb104::mTurquoise2-b104^dPAA line #2; L5, pb104::mTurquoise2-b104^dPAA line #3; L6, pUBQ10::mTurquoise2-bHLH104 line #1; L7, pUBQ10::mTurquoise2-bHLH104 line #2. (B) Representative images of 30-days-old Arabidopsis plants grown in hydroponic system (scale bar = 5 cm). Among grown plant lines, pb104::b104^dPAA #2 (L4) grew poorly and was not used in ICPMS and gene expression analysis. (C) Experimental design for the dataset shown. Arabidopsis plants were grown hydroponically under iron-sufficient (+Fe) conditions for 27 days, followed by a 3-day treatment with either +Fe or –Fe media. (D) Schematic representation of tissue separation used for both ICP-MS and RT-qPCR analysis. Young and old leaves were separated from the roots to examine tissue-specific iron accumulation and gene expression. The same tissue layout is used to represent heatmaps in the ICP-MS dataset. (E) Iron content in young leaves, old leaves, and roots under +Fe and –Fe conditions, measured by ICP-MS. Data represent mean values from three biological replicates (n = 3). Heat map intensity reflects iron concentration (µg/g dry weight). Color shading was generated using the R package ggplantmap, with deeper blue indicating higher Fe content. Asterisks indicate genotypes that differ significantly from WT within the same tissue and treatment condition (p < 0.05), as determined by two-way ANOVA and Tukey’s post hoc test (n = 3 for each group). (F) Clustered heatmaps showing the expression profiles (RNA sequencing) of selected Fe homeostasis genes in young leaves, old leaves, and roots under both +Fe and – Fe conditions. Data represent the mean of three biological replicates (n = 3). Z-scores were calculated from normalized RNA expression values and visualized in RStudio version 4.4.1. Yellow indicates high expression; purple indicates low expression.

Elemental profiling showed that *p104::b104^dPAA line #3* accumulated more Fe than control lines at the late developmental stage (Figure 4E). Under +Fe, roots of *pb104::b104^dPAA* #3 had increased Mn and Zn levels compared to WT (Supplementary Figure S9). Cu levels were similar as WT.

In transcriptome analysis, *pb104:b104^dPAA* #3 plants had the highest DEG counts in young leaves (3,877 DEGs) and old leaves (1,895 DEGs) under +Fe conditions compared with controls, while root tissue exhibited minimal differential expression differences (71 DEGs) (Supplementary Figure S10A). GO term enrichment analysis revealed that genes associated with immune response were significantly upregulated in both old and young leaves of *pb104:b104^dPAA* #3 plants under +Fe conditions. In contrast, 3-day -Fe treatment resulted in young leaves in the upregulation of terms related to Fe homeostasis (Supplementary Figure S10B). This indicates a tissue-specific transcriptional response to Fe availability at the reproductive stage.

Specifically, expression analysis showed that in roots of *pb104::b104^dPAA line #3* genes were upregulated under +Fe, similar as we reported above for the #2 line at the seedling stage, namely upregulation of *BHLHIVc* genes, Fe deficiency response, Fe sufficiency marker and negative regulator genes (Figure 4F). Interestingly, Fe deficiency response genes such as *BTS*, *IMA1*, *BHLH100*, and *PYE* were upregulated under –Fe in young leaves of pb104::b104 #1 compared to pb104::b104^dPAA line #3 (Figure 4F). *FER1* was upregulated in young and old leaves of *pb104::b104^dPAA* line #3 compared to pb104::b104 #1 and WT under +Fe. Targeted expression profiling by RT-qPCR confirmed again the transcriptome data (Supplementary Figure S11). Hence, even though at the seedling stage *pb104::b104^dPAA* line #3 did not display iron accumulation phenotypes, this was clearly the case at the flowering stage, indicating that proper bHLH104 function may be more important at later than early growth stages.

Finally, as *pb104::b104^dPAA* line #2 did not grow in +Fe hydroponic conditions, we modified the hydroponic growth system and changed to a Fe resupply assay. Seedlings were transferred at an early stage to -Fe (instead of the usual +Fe) medium for 27 days, followed by a 3-day +/- Fe resupply. Here, we found interesting phenotypes. While WT plants remained chlorotic and accumulated Fe mostly in the roots following the Fe resupply, *pb104::b104^dPAA* line #2 showed substantial Fe accumulation in root and leaves despite persistent leaf chlorosis, suggesting high root-to-shoot Fe translocation. Interestingly, *pb104::b104^dPAA* line #2 accumulated significantly less Mn, Zn, Cu compared to control plants in old and young leaves (Supplementary Figure S12). This shows that at the reproductive stage, there is a particularly elevated Fe uptake and translocation towards shoots in *pb104::b104^dPAA* #2 compared to the WT, while that of Mn, Zn and Cu is suppressed.

Since 30-day-old *pb104::b104^dPAA* #3 plants accumulated not only Fe but also Mn and Zn under + Fe in roots, we examined the possibility that this might have been caused by altered expression of metal transporter genes. We found elevated expression of metal transporter genes, including NRAMP1, COPT2, MTP8, and ZIP family members, in *pb104::b104^dPAA* roots under both +Fe and -Fe at the seedling stage, and under +Fe at the reproductive stage, consistent with the observed metal accumulation patterns (Supplementary Figure S13).

To summarize, *pb104::b104^dPAA* lines accumulated Fe throughout the tested reproductive stage. Metal accumulation and associated phenotypes due to b104^dPAA expression remained consistent from seedling to advanced developmental stages.

### The PAA short sequence is required for 26S Proteasome-mediated degradation of bHLH104

The severity of phenotypes in *pb104::b104^dPAA* lines was not explained by differences in transgene transcript levels (Supplementary Figures S8). Additionally, *pUBQ10::b104* lines showed no significant phenotype compared to WT apart from shorter roots despite elevated *bHLH104* transgene expression (Supplementary Figure S4, Supplementary Figure S8C). Therefore, we suspected that variation in bHLH104 protein accumulation underlies the observed phenotypic differences.

We thus examined *in vivo* accumulation of mTurquoise2-tagged bHLH104 and b104^dPAA protein, expressed under either the *UBQ10* or *BHLH104* promoter (Figure 5A, B). Immunoblotting of root tissue proteins detected mTurquoise2-b104^dPAA protein in *pb104::b104^dPAA* lines #1 and #2 (Figure 5B, Supplementary Figure S14). In contrast, mTurquoise2-tagged protein was barely detectable in *pb104::b104^dPAA* lines *#3* and *#4*, *pb104::bHLH104*, and *pUBQ10::b104* lines. Hence, mTurquoise2-b104^dPAA protein levels coincided with the strength of Fe accumulation phenotypes, while the wild type mTurquoise2-b104 was not detected.

**Figure 5.**
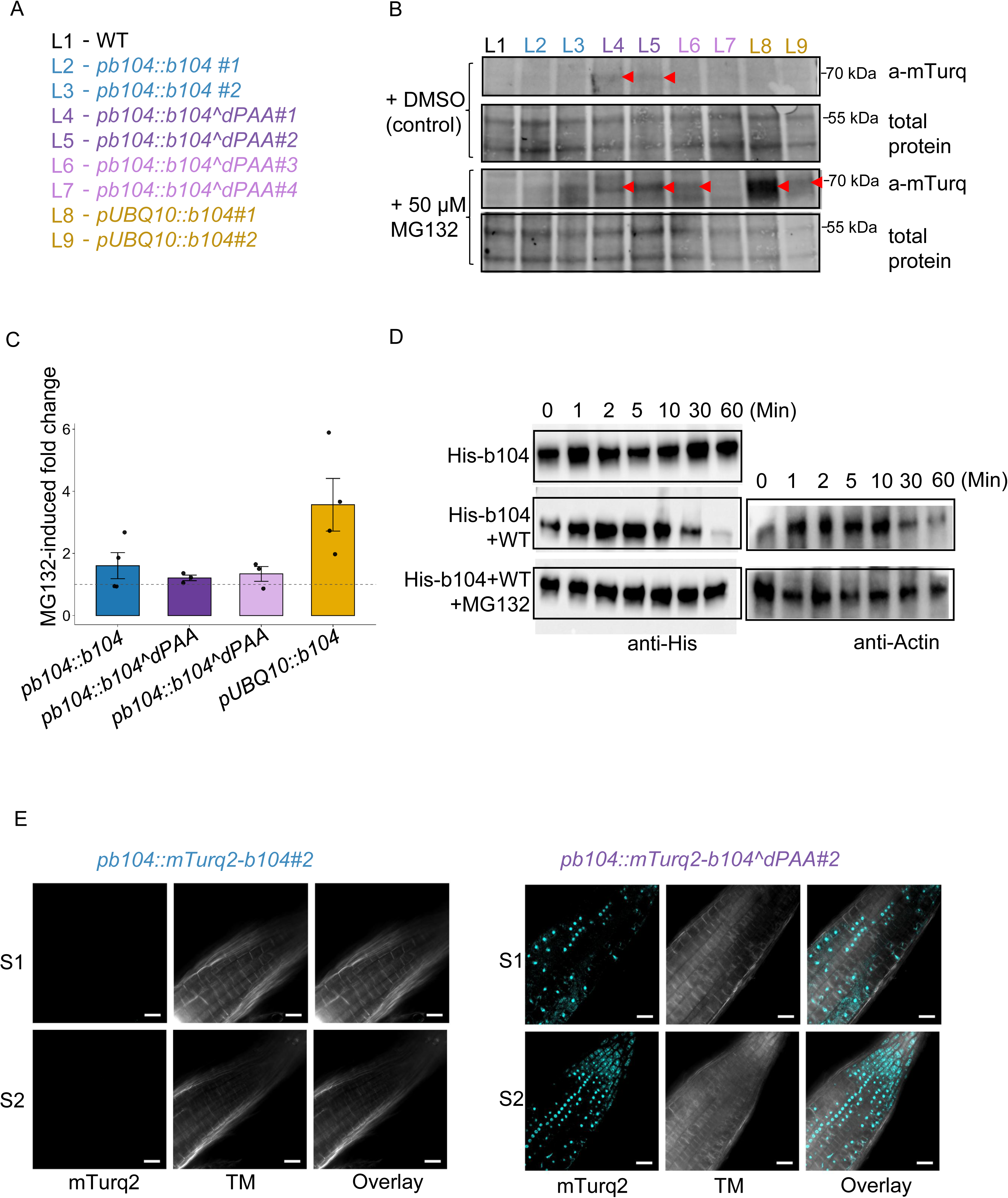
The PAA short sequence mediates bHLH104 protein stability and affects its abundance in Arabidopsis. (A) Genotypes and transgenic lines used in this experiment. (B) Immunoblot analysis of mTurquoise2-tagged bHLH104 and b104^dPAA in different transgenic lines treated with either DMSO (control) or 50 μM MG132. Total protein and anti-mTurquoise2 blots are shown. Samples are labeled from #L1 to #L9. The transgenic lines exhibiting increased protein abundance correspond to those that displayed altered growth phenotypes in Figure 1, suggesting a link between protein stability and the observed developmental responses. Red arrows indicate the expected band position for mTurquoise2-bHLH104 fusion protein. Additional blots for replicate studies are provided in Supplementary Figure S14. Original uncropped blot is shown in Supplementary Figure S19. (C) Relative protein levels of mTurquoise-tagged bHLH104 were determined by Western blot. Band intensities were normalized to total protein, and fold-change was calculated as the ratio of MG132-treated to DMSO-treated samples. Bars represent mean ± SEM; individual data points are shown. (D) Cell-free degradation assay showing in vitro degradation of His-tagged bHLH104 protein when incubated with protein extracts from 10-day-old Col-0 (WT) Arabidopsis thaliana seedlings. His-bHLH104 was expressed in E. coli and remained stable on its own, but degraded upon incubation with the plant extracts, indicating proteasome-mediated degradation. Total protein from whole seedlings was extracted using standard protein extraction procedures. Samples were collected at the indicated time points and analyzed by immunoblotting with anti-His and anti-Actin antibodies. MG132 treatment stabilizes bHLH104, indicating proteasome-dependent degradation. (E) Confocal microscopy of mTurquoise2 fluorescence in roots of transgenic Arabidopsis seedlings expressing full-length bHLH104 (left) or b104^dPAA (right). Images show two representative seedlings (S1 and S2) per genotype. Arabidopsis thaliana seedlings were at the germination stage, 5 days old, corresponding to 2 days after stratification, and grown on +Fe medium. TM, transmitted light; Overlay, merged mTurq2 and TM channels. Scale bar = 25 µm; excitation at 458 nm; emission collected from 493–613 nm.

To check whether the low abundance of mTurquoise2-bHLH104 is due to proteasomal degradation, we treated samples with proteasome inhibitor MG132. MG132 treatment stabilized full-length mTurquoise2-bHLH104 in *pUBQ10::b104* lines by more than two-fold. However, MG132 treatment did not affect the abundance of mTurquoise2-bHLH104 in any native promoter lines (Figure 5B-C). To confirm that bHLH104 is subject to proteasomal degradation, a cell-free degradation assay was performed. When recombinant His-bHLH104 protein, purified from *Escherichia coli*, was incubated with whole protein extracts from WT seedlings, it was rapidly degraded. This degradation was inhibited by the proteasome inhibitor MG132, supporting the involvement of the 26S proteasome (Figure 5D).

Confocal imaging of root tips further confirmed the increased abundance of mTurquoise2-b104^dPAA, as indicated by fluorescence signals in nuclei of root tip cells in *pb104::b104^dPAA* lines, whereas fluorescence was undetectable in lines expressing full-length mTurquoise2-bHLH104 (Figure 5E).

Altogether, these findings show that the PAA short sequence is required for degradation of bHLH104 by the 26S proteasome. Removing this short sequence seems to prevent normal turnover of the protein.

### The C-terminal PAA sequence of bHLH104 mediates interaction with BTS E3 ligases and enables ubiquitination

To understand how the PAA sequence contributes to bHLH104 degradation, we examined its role in mediating interaction with BTS.

AlphaFold2 multimer modeling (Evans et al., 2021; Jumper et al., 2021) and electrostatic surface analysis (Pettersen et al., 2004) revealed a charge-based interface with high predicted Local Distance Difference Test (pLDDT) scores, indicating strong confidence in the predicted structure (Figure 6A-B). These models suggest that the PAA sequence forms a stable and specific docking surface for BTS, likely critical for E3-substrate recognition.

**Figure 6.**
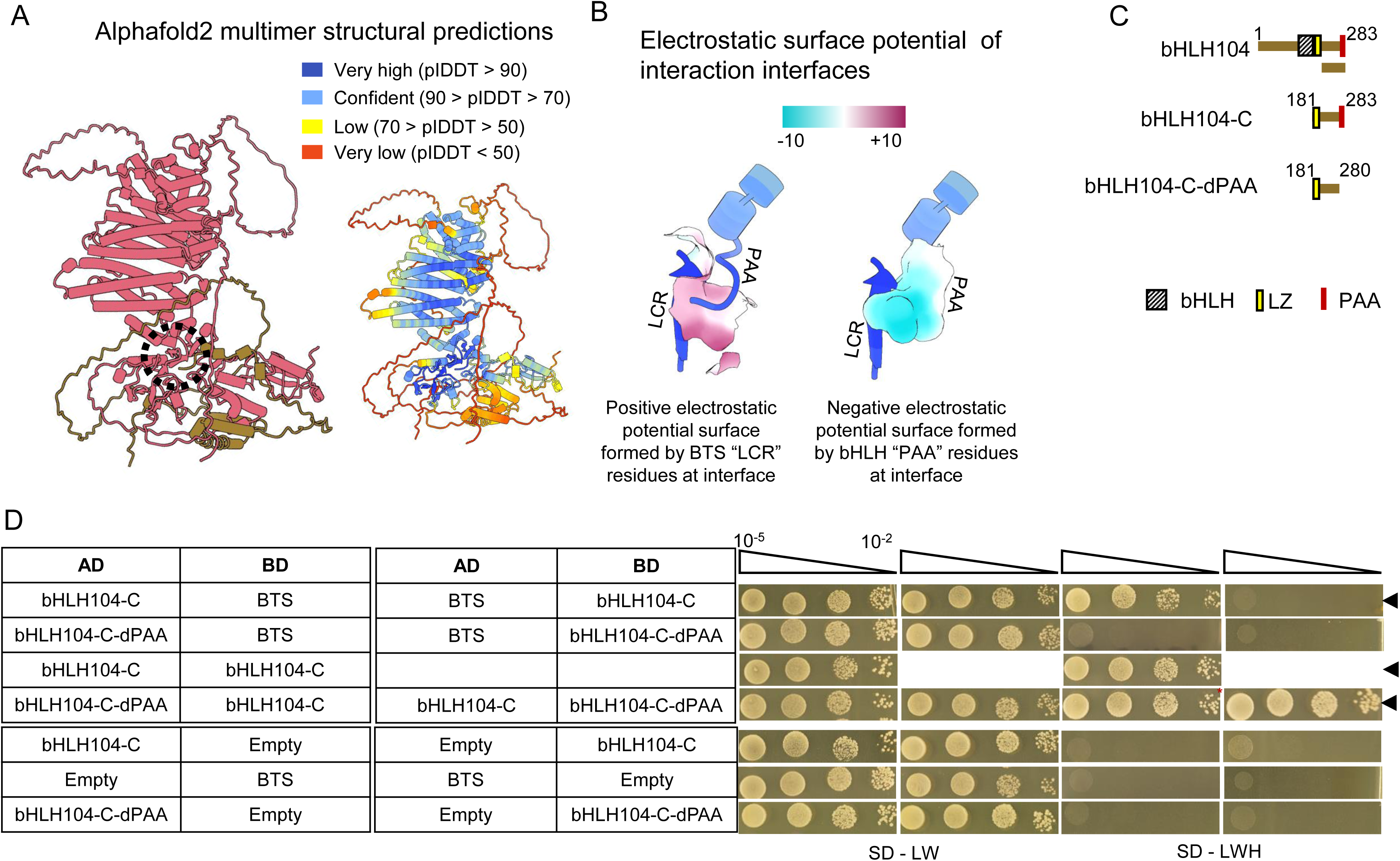
The PAA short sequence is required for bHLH104 interaction with BTS. (A) Structural prediction of the bHLH104–BTS complex using AlphaFold2 Multimer (Evans et al., 2021; Jumper et al., 2021). Left: Chain-colored model showing BTS (rose color) and bHLH104 (bronze color). Right: Per-residue confidence model with pLDDT scores color-coded as very high (pLDDT > 90), confident (90 > pLDDT > 70), low (70 > pLDDT > 50), and very low (pLDDT < 50). Dashed circle marks the interaction interface formed by BTS and bHLH104. (B) Electrostatic surface potential modeling (Pettersen et al., 2004) reveals a negatively charged region near the proline–alanine–alanine (PAA) sequence in bHLH104 that complements a positively charged surface in the leucine–cysteine–arginine (LCR) domain of BTS. The electrostatic scale ranges from –10 (purple) to +10 (cyan) kT/e. (C) Schematic representation of full-length bHLH104, bHLH104-C and bHLH104-C-dPAA used for Y2H and UbiGate assays. C-terminal parts lack the N-terminus and the DNA-binding basic Helix Loop Helix (bHLH) domain. LZ is Leuzine zipper domain. Numbers indicate amino acid positions. (D) Yeast two-hybrid assays testing protein–protein interactions between bHLH104-C^dPAA and BTS. bHLH104-C and BTS heterodimerization and bHLH104-C homodimerization serve as positive controls. Note that C-terminal bHLH104 fragments were used since full length bHLH104 shows autoactivation in Y2H (Lichtblau et al. 2022). Black arrows indicate positive interaction due to colony growth in the selective condition.

To further study protein interaction in the presence and absence of PAA experimentally, we made use of established protein expression systems. Due to the difficulties of detecting bHLH104 and BTS in plant cell assays, we opted for the yeast two hybrid (Y2H) assay (Lichtblau et al., 2022). BTS interacts non-reciprocally with the C-terminal part of bHLH104 (bHLH104-C) in yeast, and bHLH104-C does not auto-activate in the yeast-two-hybrid (Y2H) system (Figure 6C), as previously shown (Lichtblau et al., 2022). We now tested whether interaction is different with bHLH104-C^dPAA. Indeed, deletion of the PAA sequence abolished the interaction with BTS (Figure 6D). bHLH104 is known to homodimerize (Zhang et al., 2015; Li et al., 2016; Lichtblau et al., 2022). bHLH104-C^dPAA still dimerized with bHLH104-C, confirming that bHLH104-C^dPAA is present and properly folded in yeast cells. A similar loss of interaction ability was found between bHLH104-C-dPAA and BTSL1 and BTSL2 (Supplementary Figure S15), indicating that the PAA sequence is broadly required for recognition by BTS/ BTSL E3 ligases.

To test whether bHLH104 is ubiquitinated by BTS with requirement of the PAA short sequence, we established the bacterial UbiGate system (Kowarschik et al., 2018; Ortmann et al., 2023) to reconstitute a BTS-dependent ubiquitin-conjugation cascade for bHLH104 (Figure 7A). The UbiGate system co-expresses all ubiquitination and target components within living bacterial cells, providing a cellular context for a direct ubiquitination reaction while enabling unambiguous assignment of E3-substrate specificity through distinct epitope tags on each component (Figure 7B, C). Differences in banding patterns between controls lacking individual cascade components and cells containing the complete cascade are indicative of ubiquitination. Since full-length BTS was not stably expressed in *E. coli*, we used a truncated version (BTS-HC) lacking the first two hemerythrin domains. The relevant assay consists of comparing the Strep-tagged bHLH104-C protein patterns in the presence (lanes 1, 2, and 5 of Figure 7B,C) and absence of E2 (UBC8) (lanes 3 and 6 of Figure 7B,C), and E3 (BTS-HC) lanes 4 and 7 of Figure 7B,C). Immunodetection with the anti-Strep antibody confirmed that bHLH104-C was expressed in all sample cultures, with a band at the expected size of the unmodified protein (∼40 kDa), as indicated by brown arrows in Figure 7B. Some higher molecular weight bands occur independently of the E3 ligase and are therefore not specific to BTS-HC activity, as indicated by grey arrows in Figure 7B. However, in the presence of all cascade components, one band (∼70 kDa),as indicated by a green arrow in Figure 7B, appears that is absent from b104-C^dPAA under identical conditions and absent from the substrate-negative control (Figure 7B, Supplementary Figure S16). The size of this band is consistent with polyubiquitination of bHLH104-C, as its molecular weight shift of approximately 30 kDa corresponds to the addition of three to four ubiquitin moieties, resembling a characteristic ubiquitination pattern (Kowarschik et al., 2018). This meets the minimum chain length of four ubiquitins required for efficient 26S proteasome recognition (Thrower, 2000). These data suggest that the PAA short sequence is required for BTS-mediated ubiquitination of bHLH104. BTS autoubiquitination, reported previously (Kobayashi et al., 2013; Selote et al., 2015), served as a positive control for the ubiquitination reaction in the UbiGate system (Figure 7C). bHLH115 has been previously indicated as a target of BTS-dependent degradation (Selote et al., 2015; Xing et al., 2021). Consistent with these reports, we found that bHLH115 is also likely ubiquitinated by BTS-HC in the UbiGate system, indicating that this assay is suited (Supplementary Figure S17).

**Figure 7.**
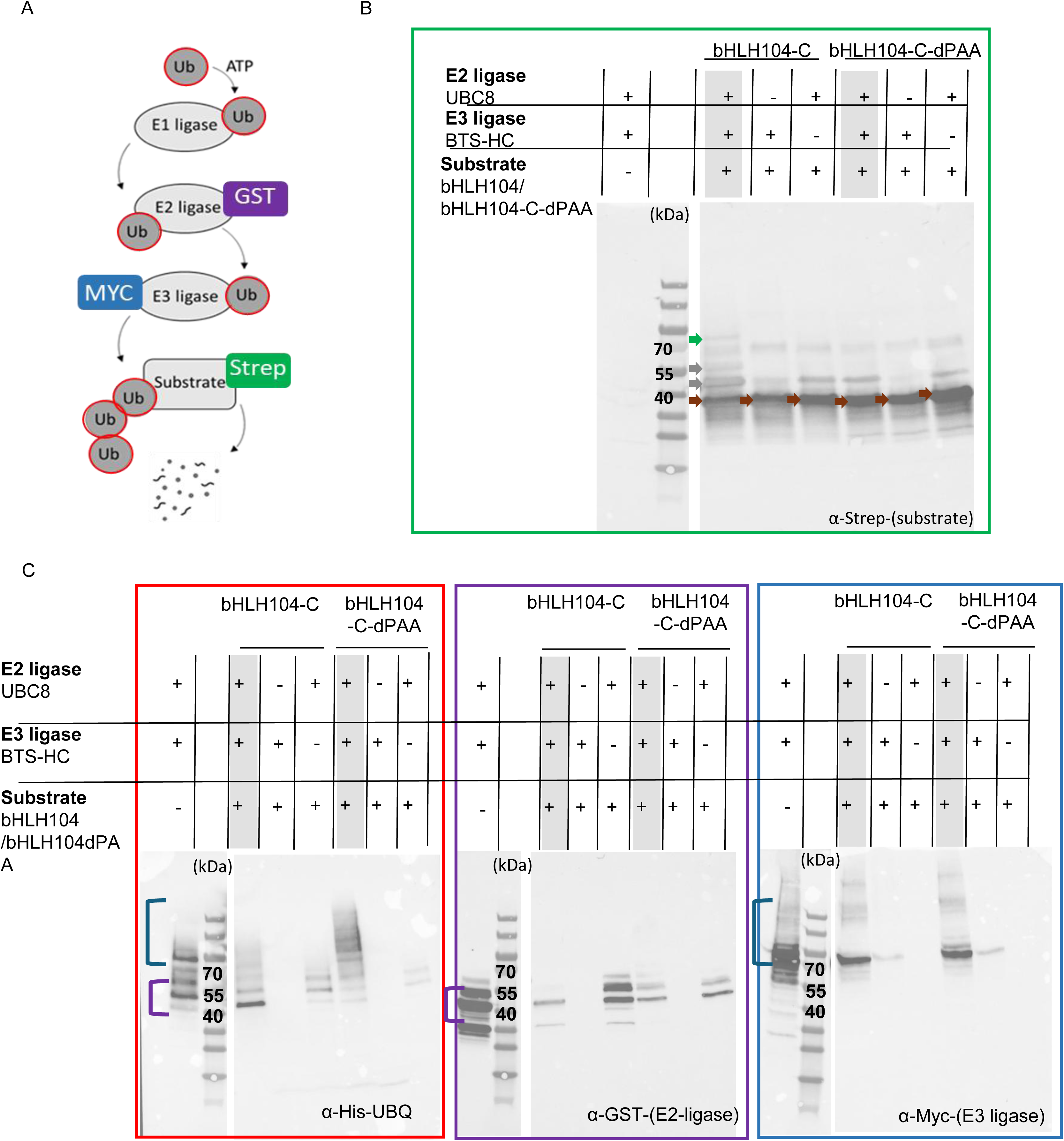
The PAA short sequence leads to a specific bHLH104-C band in the UbiGate system testing for BTS-mediated ubiquitination of bHLH104 in E. coli cells. (A) Schematic overview of the ubiquitin/26S proteasome pathway. Each component was expressed in E. coli using the UbiGate system (Kowarschik et al., 2018). Recombinant bHLH104-C or bHLH104-C^dPAA proteins (Strep-tagged) were used as substrates. BTS-HC served as the E3 ligase (Myc-tagged), in combination with UBC8 as E2 ligase (GST-tagged), an E1 activating enzyme, and ubiquitin (His-tagged). (B) The relevant immunoblot for assessing substrate ubiquitination is the anti-Strep blot. The anti-Strep antibody detects predominantly the Strep-tagged substrate protein in as an expected band corresponding to a non-ubiquitinated form (∼40 kDa; brown arrows). Only in the presence of all components, there are multiple higher than 40 kDa protein bands, indicative of a protein ubiquitination ladder (Kowarschik et al., 2018), with one particular unique form (∼70 kDa) potentially corresponding to polyubiquitinated bHLH104-C (green arrow corresponding to ∼70 kDa plus molecular weight shift of approximately 30 kDa corresponding to the addition of three to four ubiquitin moieties of ∼8.6 kDa each; grey arrows indicating additional higher molecular weight bands indicating a ubiquitination ladder pattern). The 70 kDa band is absent in the relevant controls, either bHLH104-C^dPAA lane under identical conditions or in control samples lacking individual components. Strep bands are also detectable in control lanes, these are not considered ubiquitination signals. The assay has been performed three times with similar outcome of the unique 70 kDa band which might point to the polyubiquitinated bHLH104-C target (additional replicates in Supplementary Figure S16). (C) Shown are three types of control immunoblots confirming expression of all proteins for ubiquitination activity of the E3 ligase in the UbiGate assay. All panels are from the same membrane, sequentially probed with different antibodies after stripping. Anti-GST detects E2 ligase (GST-UBC8); anti-Myc detects E3 ligase (Myc-BTS-HC); anti-His-UBQ detects total ubiquitin conjugates (His-Ub). Violet bracket indicates autoubiquitinated E2 ligase; blue bracket indicates autoubiquitinated E3 ligase. Bands in control lanes of the anti-ubiquitin blot represent E2 and E3 autoubiquitination, which is independent of substrate ubiquitination.

These results suggest that bHLH104 can act as a substrate of BTS in a cellular environment and that the PAA short sequence is required for both physical interaction and ubiquitination. The specificity of this finding is supported by the bHLH104-C^dPAA control, which differs from bHLH104-C by only three amino acids yet does not show the specific band, indicating that the presence of the PAA short sequence is required for interaction and ubiquitination by BTS.

## Discussion

This study demonstrates functional relevance of the terminal PAA short sequence of bHLH104 by studying a PAA deletion variant. Our data showed and are interpreted as follows, in agreement with recent literature on other bHLHIVc transcription factors: bHLH104 is degraded via the proteasome. Lack of PAA prevents binding of bHLH104^dPAA to BTS E3 ligase. Hence, bHLH104^dPAA is not ubiquitinated nor targeted by the proteasome. This leads to protein stabilization and the observed higher abundance of bHLH104^dPAA. bHLH104^dPAA is functionally active. Its enhanced abundance causes target genes of the Fe deficiency response pathway to be constitutively up-regulated throughout plant development. Resulting excessive uptake of Fe and misregulation of Fe homeostasis lead to accumulation of Fe and other metals. This causes oxidative stress and perturbs plant performance, resulting in small plants exhibiting Fe toxicity in leaves. Yet, the level of bHLH104^dPAA protein can vary in transgenic lines, so that some lines can accumulate Fe over their life span but their growth is not compromised.

### bHLH104 abundance is controlled via the proteasome

Our results show that wild-type bHLH104 protein is susceptible to degradation in a plant cell environment, and inhibition of the proteasome pathway increases bHLH104 abundance. The post-translational regulation of bHLH IVc proteins has been previously reported, with transcript levels remaining largely unchanged under Fe deficiency, suggesting that regulation occurs primarily at the protein level (Zhang et al., 2015; Li et al., 2016; Liang et al., 2017; Selote et al., 2015). Selote et al. (2015) showed in a cell-free degradation assay that bHLH105/ILR3 and bHLH115 were degraded when mixed with wild-type seedling whole protein extracts, whereas bHLH104 remained stable even after 8 hours of incubation. In contrast, our recombinant bHLH104 purified from *E. coli* was degraded within 1 hour in a proteasome-dependent manner (Figure 5D). This difference may be attributed to the protein expression system, as Selote et al. (2015) used in vitro translated protein from wheat germ extract, which may differ from bacterially expressed protein in folding or modifications that affect susceptibility to degradation. Proteasomal degradation of bHLH104 is further supported by our in planta observations: mTurquoise2-bHLH104 protein was barely detectable by immunoblotting in transgenic lines, including *pUBQ10::b104* lines despite their high transcript levels, and MG132 treatment resulted in increased protein detection (Figure 5B, C). Consistently, mTurquoise2 fluorescence was undetectable by confocal imaging in lines expressing full-length bHLH104, whereas b104^dPAA lines showed clear nuclear signals (Figure 5E). Our data are consistent with the overall model of proteasomal regulation of bHLH IVc proteins and extend it to bHLH104 specifically, as our UbiGate data indicate that BTS can ubiquitinate bHLH104 in a PAA-dependent manner. In line with this, a recent study showed that BTSL1 and BTSL2 also target bHLH105 and bHLH115 for degradation in roots (Zhao et al., 2026), indicating that multiple BTS-type E3 ligases act redundantly on bHLH IVc proteins in root tissue, where Fe acquisition takes place. Interestingly, MG132 treatment also resulted in higher b104^dPAA abundance (Figure 5B, C), even though b104^dPAA escapes BTS-mediated ubiquitination. This indicates that PAA-independent proteasomal degradation mechanisms also contribute to bHLH104 turnover. Such alternative degradation could involve additional E3 ligase systems, as demonstrated for the apple orthologue MdbHLH104, which is targeted by the CUL3-BTB/TAZ complex MdBT2-MdCUL3 independently of HRZ/BTS (Zhao et al., 2016), or could involve selective autophagy, which has been shown to degrade transcription factors in plants (Wang et al., 2021; Raffeiner et al., 2023). This would explain why complete protein stabilization is not achieved by PAA deletion alone.

Interestingly, despite elevated *mTurq2-bHLH104* transcript levels in shoots, *pUBQ10::b104* lines showed no phenotype, in contrast to previously reported *p35S::bHLH104-GFP* lines that displayed enhanced Fe deficiency tolerance, increased ferric-chelate reductase activity, and Fe overaccumulation (Zhang et al., 2015). Several factors may account for this discrepancy. The 35S promoter typically drives stronger expression than the *UBQ10* promoter, potentially saturating BTS-mediated degradation capacity. Additionally, the C-terminal GFP tag in *35S::bHLH104-GFP* lines may affect BTS accessibility to the adjacent PAA short sequence, whereas an N-terminal mTurquoise2 tag leaves the C-terminus unobstructed. The *pb104::b104^*dPAA line #2 exhibited upregulation of *ILR3/BHLH105, BHLH115, BHLH39* and *BHLH100* in roots under both +Fe and -Fe conditions. This indicates that functional output of bHLH104 depends on protein accumulation rather than transcript abundance. Whether the observed upregulation of *BHLHIVC* genes reflects direct transcriptional autoregulation or indirect effects through altered Fe status remains to be determined; however, these findings suggest a potential positive feedback loop within the bHLH IVc network that is gated by BTS-mediated degradation. The direct interaction of bHLH104 with the BTS E3 ligase has been reported previously (Long et al., 2010; Selote et al., 2015; Lichtblau et al., 2022), and genetic evidence supported a regulatory upstream relationship of BTS towards bHLH104 (Zhang et al., 2015). We tested this regulatory relationship and identified that the PAA short sequence is needed for BTS interaction with bHLH104. We propose a model (Figure 8) in which bHLH104 protein is maintained at low abundance through proteasome-mediated degradation. A limitation of this study is the difficulty in visualizing mTurquoise2-tagged full-length bHLH104 in planta due to rapid degradation, whereas b104^dPAA was detectable in early developing seedlings (Figure 5E). Future studies should investigate additional post-translational modifications and identify environmental and physiological conditions that promote bHLH104 stabilization.

**Figure 8.**
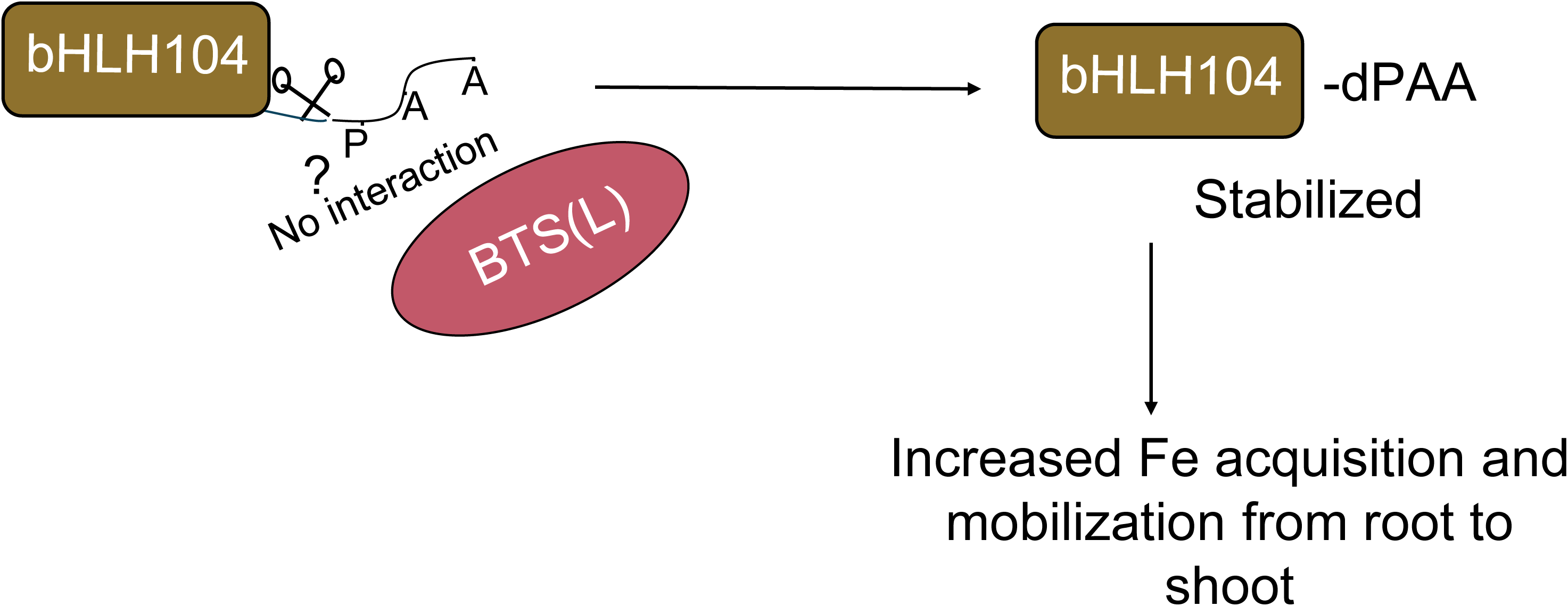
Working model of bHLH104 regulation through the C-terminal PAA sequence and its impact on Fe homeostasis. The C-terminal PAA sequence in bHLH104 mediates interaction with BTS-type E3 ligases, targeting the protein for proteasomal degradation. Deletion of the PAA motif (b104^dPAA) disrupts this interaction, leading to protein stabilization. The increased abundance of stabilized b104^dPAA enhances Fe acquisition responses in Arabidopsis.

### b104^dPAA stimulates Fe and metal ion accumulation and potential for biofortification

Multiple metals accumulated in *pb104::b104^*dPAA plants. The multi-metal accumulation can be explained by persistent activation of a broad array of metal transporters in root cells, like Fe-, Zn and Mn-transporting IRT1, Fe- and Mn-transporting NRAMP1, Cu-transporting COPT2, Mn-transporting MTP8 (Connolly et al., 2002; Vert et al., 2002; Cailliatte et al., 2010; Eroglu et al., 2016). The genes encoding these transporters are under control of the FIT transcription factor (Schwarz and Bauer, 2020), and the FIT pathway was found active downstream of bHLH104 in our study. Since b104^dPAA plants accumulate excess metals including Zn, this could further amplify the Fe deficiency signal. This is consistent with reports that heavy metals like Zn can induce Fe deficiency responses at the level of Fe sensing rather than by directly inhibiting Fe transport (Lešková et al., 2017), and with the demonstration that Zn or Fe binding to multiple domains of HRZ E3 ligases modulates their stability and function in response to Fe nutritional status (Shinkawa et al., 2025). Thus, the accumulated metals in *pb104::b104^dPAA* plants may reinforce Fe deficiency signalling, creating a positive feedback loop that sustains constitutive Fe uptake. Similar phenotypes where Fe accumulation disrupts plant growth have been reported in *BTS* loss-of-function mutants and in overexpression lines of *BHLH39* or *BHLH105* (Zhang et al., 2015; Li et al., 2016; Hindt et al., 2017; Naranjo-Arcos et al., 2017). A still unresolved question is why the known negative post-translational regulation mechanisms, e.g. IRT1 ubiquitination and degradation under low Fe and excessive Zn and Mn, are not in place here (Dubeaux et al., 2018; Abuzeineh et al., 2022). Perhaps this type of regulation requires that Zn and Mn are supplied in the rhizosphere rather than inside the root. In our experimental settings, excessive Zn and Mn were not supplied and Zn and Mn accumulated due to plant internal imbalance.

We observed that b104^dPAA protein abundance in seedling roots correlated with phenotype severity among *pb104::b104^dPAA* lines. A plausible explanation is that this variation in protein abundance across lines results from differences in transgene copy number arising from the floral dip transformation method (Gelvin, 2003; Nagaya et al., 2005; Meiser et al., 2011; Li et al., 2016). Notably, *pb104::b104^dPAA line #3*, which showed negligible b104^dPAA protein in seedling roots, grew normally during early development but accumulated Fe in leaves at the reproductive stage. This suggests that minimal b104^dPAA activity can increase Fe content without causing severe growth defects, highlighting potential for Fe biofortification strategies. We acknowledge that our transgenic lines have b104^dPAA along with endogenous bHLH104 protein activity, so that b104^dPAA alone or together with endogenous bHLH104 could contribute to the observed phenotypes. Additional investigations will be required to define the precise threshold at which bHLH104^dPAA protein levels enhance Fe accumulation without compromising plant growth. As demonstrated by the growth defects in strong b104^dPAA -expressing lines, careful optimization of protein levels would be required to achieve Fe biofortification without compromising plant health. The milder phenotype of *pb104::b104^dPAA line #3*, which showed elevated leaf Fe content at reproductive stage without severe growth defects, suggests that fine-tuning of this system may be achievable through careful selection of lines with appropriate expression levels. More broadly, the findings integrate with emerging work on BTS/BTSL hemerythrin-based Fe/oxygen sensing (Selote et al., 2015; Pullin et al., 2025) and clarify how post-translational control translates Fe status into transcriptional outputs.

### Pleiotropic and tissue-specific effects of b104^dPAA

The phenotypes observed in *pb104::b104^dPAA* plants extend beyond simple Fe accumulation, suggesting that constitutive Fe deficiency signaling has broader physiological consequences. Several phenotypes observed in these lines may reflect activation of multiple stress regulatory pathways. First, the delayed flowering phenotype may result from -Fe response signaling interfering with flowering time control through epigenetic, circadian, and hormone pathways (Cheng et al., 2004; Salomé et al., 2012; Chen et al., 2013; Hong et al., 2013; Wild et al., 2016; Dutta et al., 2017; Maruoka et al., 2022). Second, gene ontology analysis of *pb104::b104^dPAA* leaves showed enrichment of immune signaling and salicylic acid signaling terms. Fe deficiency treatment is known to elevate salicylic acid levels and induce salicylic acid-responsive genes (Shen et al., 2016), and bHLH105 can integrate Fe homeostasis with plant immune responses against pathogens (Rampey et al., 2006; Aparicio and Pallás, 2017; Samira et al., 2018). This suggests that the Fe deficiency response is interconnected with immune signaling pathways.

Expression of b104^dPAA in *pb104::b104^dPAA* lines results in distinct tissue-specific consequences. The lack of upregulation of internal Fe allocation genes such as *BTS, PYE, IMA1*, *BHLH100, BHLH39* under -Fe condition in the leaves of mature *pb104::b104^dPAA* lines suggests that elevated local Fe levels in the leaves suppress the Fe deficiency response downstream of bHLH104, consistent with recent evidence that leaf Fe responses are determined mainly by local Fe status rather than root-derived systemic signals (Nguyen et al., 2022; Ngigi et al., 2025). In contrast, in roots of *pb104::b104^dPAA* lines, -Fe acquisition genes such as *FIT*, *IRT1* and *FRO2* were upregulated. This indicates that, rather than systemic signaling, an active and constitutive Fe acquisition response is maintained in the roots. In wild-type plants, bHLH104 is expected to be stabilized only under Fe deficiency, enabling transient activation of uptake genes. Lack of PAA prevents this regulation, resulting in constitutive bHLH104 activity and continuous Fe uptake regardless of Fe status. Interestingly, we observed enhanced Fe mobilization to young leaves in *pb104::b104^dPAA line #2* after 3 days of Fe supply following growth in –Fe. In contrast, wild-type plants retained most Fe in roots The enhanced root-to-shoot Fe mobilization in pb104::b104^dPAA line #2 may be partly explained by the elevated IRT1 expression in these lines. (Quintana et al., 2022) showed that IRT1 protein, independent of its Fe(II) transport activity, is necessary for effective root-to-shoot Fe partitioning, suggesting that constitutive IRT1 upregulation driven by b104^dPAA could promote Fe translocation to shoots.. This shows that bHLH104 protein stability serves as a critical control point linking Fe sensing to transcriptional responses in roots, whereas leaves respond primarily to local Fe availability through regulatory mechanisms independent of bHLH104 protein stability. Grafting experiments would help determine whether the effects of stabilized bHLH104 are root- and leaf-autonomous or involve systemic signaling.

### The PAA short sequence is critical for protein regulation

The present study found that the terminal PAA sequence of bHLH104 mediates critical interactions. Additionally, small proteins IMA1 and IMA3, which interact with BTS and BTSL1, also contain a conserved C-terminal PAA sequence, indicating that this sequence is broadly conserved for E3 ligase recognition (Li et al., 2021; Lichtblau et al., 2022). Other members of the bHLH IVc family have instead a PVA terminal sequence. It has not yet been examined which role PVA plays and whether it is similar to PAA. One possible explanation is that the middle alanine in the PAA sequence of bHLH104 may not be essential. Previous studies have shown the significance of terminal alanine residue in bHLH IVc protein regulation. The terminal alanine in bHLH115 and bHLH105 is essential for BTS interaction (Li et al., 2021), while overexpression of bHLH34 with its terminal alanine substituted to valine resulted in a metal accumulation phenotype similar to our observations (Sharma and Yeh, 2020). Taken together, these observations support the conclusion that the PAA sequence is required for the recognition of bHLH104 by the proteasomal degradation system. Our findings refine the BTS recognition requirement from the larger bHLH104 C-terminal deletions previously used to map this interaction to a three-residue sequence. The conservation of this sequence in bHLH104 orthologues, and its presence with minor variations in other bHLH IVc members, corrobates the evidences on the conserved BTS(L) recognition interface within the family. In future studies, the specificity and position effects of each amino acid in the PAA and PVA short sequences can be further examined. It will also be interesting to test whether artificial addition of PAA to transcription factors can target them for degradation by BTS/L proteins under Fe deficiency, which would establish PAA as both necessary and sufficient for BTS recognition. If so, this can be exploited in future synthetic and plant biotechnological approaches.

### Concluding remarks and perspectives

This study provides evidence that a short three–amino acid PAA sequence in bHLH104 is required for post-translational regulation of protein abundance and activity.

The conserved PAA sequence provides opportunities for biotechnological applications. As bHLH IVc proteins are conserved across plant species (Jiang et al., 2024), genome editing of this sequence in crop orthologues may enhance Fe accumulation for biofortification. In contrast to whole-gene overexpression, precise PAA editing offers potential to increase Fe content while minimizing pleiotropic effects. However, because the PAA sequence is also present in IMA proteins, which positively regulate Fe uptake, editing approaches must specifically target bHLH IVc genes without affecting *IMA* genes. Testing whether PAA addition to other transcription factors targets them for BTS/BTSL-mediated degradation could enable Fe-responsive protein degradation systems for synthetic biology applications.

Overall, our findings provide comprehensive evidence which shows that post-translational regulation of bHLH104 is necessary to maintain a balance between Fe acquisition and normal growth and development, and that disrupting this control can lead to wide-ranging physiological effects. This research has direct impact for improving crop productivity on alkaline/calcareous soils and under climate stress. Identifying sequence motifs that control master regulators of Fe uptake such as bHLH104 provides paths for crop improvement and biofortification. Our finding is that a three–residue PAA short sequence in bHLH104 can be a precise target for tuning Fe acquisition in plants. Given current efforts to enhance micronutrient use efficiency and nutritional quality, these insights are both mechanistically fundamental and directly translatable.

## Materials and methods

### Gene Accession numbers

The following *Arabidopsis thaliana* genes were the focus of this study: bHLH104 (AT4G14410), BTS/BRUTUS (AT3G18290), BTSL1 (AT1G74770), and BTSL2 (AT1G18910).

### Plant Material and Growth Conditions

*Arabidopsis thaliana* Columbia-0 (Col-0) was used as the wild type in this study. The transgenic lines were generated using the floral dip method using Rhizobium radiobacter strain GV3103 (*Agrobacterium tumifaciens)* (Clough and Bent, 1998). Constructs for generating transgenic lines were assembled using the GreenGate cloning system (Lampropoulos et al., 2013). The following modules were assembled into pGGZ003 destination vector: Promoter sequence (pUBQ10/ pbHLH104 (1636-bp sequence upstream of *BHLH104* gene)), an N-terminal mTurquoise2 fluorescent tag, Coding sequence (*BHLH104/BHLH104^dPAA*), bHLH104 terminator sequence (975 bp), Hygromycin selection marker. Primers used are listed in Supplementary Table 1.

For plate-based experiments, plants were grown as described in (Lingam et al., 2011; Lichtblau et al., 2022) in modified half-strength Hougland medium. Fe was supplied as 50 µM Fe(III)-EDTA for +Fe or omitted for -Fe conditions. The efficacy of Fe deficiency treatment was validated by confirming upregulation of canonical Fe deficiency marker genes (*IRT1, FRO2, bHLH39, bHLH100*) in WT plants (Supplementary Figure S18).

All plants for the phenotyping experiments were genotyped for the presence of transgene. For hydroponic experiments, seedlings were initially grown on modified half-strength Hoagland agar plates for 12 days before being transferred to a hydroponic setup as described in (Ngigi et al., 2025).

For soil-based phenotyping experiments, seedlings were transferred to soil on the eighth day after germination. Phenotypic measurements were taken after three weeks of soil growth.

### Phenotyping and Image Acquisition

Automated phenotyping was performed using the MicroScan system equipped with a PlantEye F600 sensor (Phenospex, Heerlen, The Netherlands) and controlled by the implemented software HortControl (Phena version 2.0, HortControl version 3.85) as previously described (Arend et al., 2016; Hüther et al., 2020; Knopf and Bauer, 2025). The PlantEye system recorded three-dimensional and spectral parameters. Spectral data included, the normalized difference vegetation index (NDVI), calculated as (near-infrared reflectance [NIR] − red reflectance [RED])/(NIR + RED).

Manual morphological phenotyping like flowering time, root length was performed as described in (Knopf and Bauer, 2025).

Fe reductase activity was measured using a protocol described in (Gratz et al., 2019). Each assay included at least three biological replicates.

Seed number was estimated based on seed weight. For each biological replicate, 100 seeds were manually counted and weighed to determine the mean weight per 100 seeds. The total seed number was calculated as (total seed weight/ weight of 100 seeds) × 100. Three biological replicates were analyzed for each genotype.

### Plant Protein Extraction and Immunoblotting

Total proteins were extracted from roots of Arabidopsis thaliana seedlings grown in modified half-strength Hougland media plates. Proteins were solubilized in 2× SDG buffer (125 mM Tris-HCl, pH 6.8, 5% SDS, 4% DTT, 20% glycerol, 0.02% bromophenol blue) at a ratio of 10 µl buffer per 5 mg homogenized root tissue. After rotation for 10 minutes and centrifugation (13,000 rpm, 10 minutes, 4°C), supernatants were collected and boiled at 60°C for 10 minutes.

Proteins were separated by SDS-PAGE using 4–20% Mini-PROTEAN TGX Stain-Free gels (Bio-Rad) and transferred to 0.45 µm nitrocellulose membranes (Bio-Rad) with the Trans-Blot Turbo Transfer System (Bio-Rad).

Membranes were blocked for 5 minutes in EveryBlot Blocking Buffer (Bio-Rad) and incubated for 1 hour at room temperature with primary antibodies against mTurquoise2 (Biozol, NBS-ASJ-L9MMEL-60; 1:1000 dilution) in 2.5% TBST with blocking buffer. After three washes in 1× TBST, membranes were incubated with HRP-conjugated secondary antibody - mouse anti-goat for mTurquoise2 [Invitrogen, G-21040; 1:1000 dilution] in TBST with 5% milk] for 1 hour. Chemiluminescent signals were developed with ECL substrate (Amersham ECL Select or SuperSignal West Femto) and imaged using the ChemiDoc MP Imaging System.

To assess the effect of proteasome inhibition on bHLH104 protein stability, 10-day-old Arabidopsis seedlings were transferred to liquid Hoagland medium and incubated for 18 hours in a plant growth chamber under the same growth conditions described above. Seedlings were treated with either 50 µM MG132 or an equal volume of DMSO (control). After treatment, total protein from roots was extracted and analyzed by immunoblotting. Band intensities were quantified using Biorad-Image Lab™ Software, and normalized to total protein staining. For each genotype, three independent biological replicates were used.

### Cell-Free Protein Degradation Assay

The coding sequence of bHLH104 was cloned into an expression vector containing an N-terminal His tag and transformed into *Escherichia coli Rosetta (DE3)* cells. Single colonies were cultured overnight in LB medium with ampicillin, and secondary cultures were inoculated at 1% volume, grown at 37°C to OD₆₀₀ = 0.6–0.8, and induced with 1 mM IPTG at 28°C for 10–12 hours. Cells were harvested by centrifugation and stored at –70°C.

For purification, frozen cell pellets were thawed on ice, resuspended in lysis buffer (30 mM Tris-HCl, pH 8.0, 150 mM NaCl) supplemented with protease inhibitors (Roche), and sonicated. Lysates were clarified by centrifugation (14,000 rpm, 30 minutes, 4°C), and His-tagged bHLH104 protein was purified using a Ni-NTA column (Cytiva) on a Bio-Rad NGC chromatography system. Bound proteins were batch-eluted with 250 mM imidazole in lysis buffer, and fractions containing the target protein were pooled, concentrated using Amicon centrifugal filters, and quantified using absorbance at 280 nm with a NanoQuant plate (Tecan). Protein concentrations were calculated using the molar extinction coefficient predicted by ExPASy ProtParam, and proteins were aliquoted and stored at –70°C.

We followed protocol from Wang et al., 2009, for cell-free degradation assay. Degradation reactions were initiated by incubating 50 µl of plant extract with purified His-bHLH104 protein at 22°C. Protein extracts were treated with either 160 µM MG132 or an equal volume of DMSO (control). Reactions were stopped at indicated time points by adding SDS loading buffer and boiling at 95°C for 10 minutes. Proteins were separated by SDS-PAGE, transferred to nitrocellulose membranes, and detected by immunoblotting using an HRP-conjugated anti-His antibody (Thermo Fisher Scientific, MA1-21315-HRP).

### Confocal Microscopy

Confocal imaging was performed using a Zeiss LSM 880 laser-scanning confocal microscope equipped with a 40× C-Apochromat water immersion objective. Two-day-old Arabidopsis thaliana seedlings grown under +Fe conditions were used for imaging. Excitation of mTurquoise2 fluorescence was carried out using a 458 nm laser line, and emitted fluorescence was collected between 493 nm and 613 nm. Images were processed and analyzed using the manufacturer’s software ZEN lite (Zeiss).

### ICP-MS Analysis

Tissues were harvested, washed sequentially in Tris-EDTA buffer and deionized water, dried on tissue paper, transferred into pre-weighed 5-ml Falcon tubes, and oven-dried at 65°C for up to two weeks. Tissue digestion was performed with 67% nitric acid (ICP-MS grade) at volumes adjusted to sample mass (1 ml for 20–35 mg, 500 µl for 10–20 mg, 350 µl for <10 mg). Samples were incubated at room temperature overnight and then heated at 95°C until clear. After centrifugation (4000 rpm, 30 minutes, 4°C), supernatants were diluted with deionized water to maintain a final acid concentration below 5% (v/v). Elemental analysis was performed using inductively coupled plasma-mass spectroscopy (ICP-MS, Agilent 7700). ICP-MS measurements were corrected for background signal and normalized by sample dilution factors.

### RNA Isolation and RT-qPCR Analysis

Total RNA was extracted using the RNeasy Plant Mini Kit (Qiagen) with on-column DNase digestion. First-strand cDNA synthesis was performed with the RevertAid Reverse Transcriptase kit (Thermo Fisher Scientific) using 300–500 ng RNA per reaction. RNA was treated with DNase I, followed by heat inactivation with EDTA and oligo(dT) primer annealing. Reverse transcription was carried out at 42°C for 1 hour. cDNA was diluted 1:10 and 1:100 for downstream applications and stored at –20°C.

Quantitative PCR (qPCR) was performed according to protocol in (Ngigi and Bauer, 2023). Each sample was measured in technical duplicates alongside a six-point standard curve (10⁷–10² copies) for each gene. The PCR program included an initial denaturation (95°C, 3 min) followed by 40 cycles of 95°C denaturation (10 sec), annealing at gene-specific temperature (15 sec), and extension (5 sec). Melt curve analysis was conducted from 65°C to 95°C.

Starting quantities were calculated from the standard curves and normalized as described in (Ngigi and Bauer, 2023; Ngigi et al., 2025). Normalized expression values were visualized as z-score heatmaps using R.

### RNA Sequencing and analysis

RNA sequencing (RNA-seq) was performed by BMKGENE (Biomarker Technologies). Library preparation and sequencing were conducted according to the service provider’s standard protocols. Data was processed following protocol in (Mai, 2024).

For DEG analysis, RNA-seq matrices were analyzed in R (v4.3+) with limma (v3.58+) using the limma-trend workflow appropriate for TPM data. Expression values were log2-transformed as log2(x + 0.5), where x is TPM value. For each comparison (line vs WT), linear models were fit with empirical Bayes moderation, and contrasts were estimated to obtain log2 fold-changes (log2FC) and Benjamini–Hochberg (BH)-adjusted P values. Genes were called differentially expressed if FDR ≤ 0.05 and |log2FC| ≥ 0.58 (∼1.5-fold).

### Gene Ontology enrichment

GO Biological Process enrichment was performed with clusterProfiler (v4.10+) against the appropriate org.*.eg.db annotation. Up- and down-regulated DEG sets were tested separately. The gene universe for each test comprised all genes assessed in the corresponding comparison. Over-representation p values were computed by the hypergeometric test and adjusted by Benjamini–Hochberg; terms with FDR ≤ 0.05 were considered significant. When indicated, redundant terms were condensed with simplify() (Wang similarity measure (Wang et al., 2007); cutoff = 0.7), retaining the most significant representative.

### Bioinformatic analyses

Protein domain architecture of bHLH104 was analyzed using the MyHits Motif Scan server (https://myhits.sib.swiss/cgi-bin/motif_scan) (Pagni et al., 2007). bHLH domain was identified at residues 121-181. The leucine zipper (LZ) domain was identified at residues 181-202. Protein structures for bHLH104, BTS and its interaction interfaces were predicted using AlphaFold Multimer (Evans et al., 2021; Jumper et al., 2021). For each protein-protein complex, 25 models were generated (5 models × 5 random seeds) to ensure comprehensive conformational sampling. Model quality was assessed using pLDDT (predicted Local Distance Difference Test) scores for per-residue confidence and ipTM (interface predicted Template Modeling) scores for interface accuracy.

The resulting structures were visualized and analyzed using UCSF ChimeraX (Pettersen et al., 2004).

### Yeast Two-Hybrid Assay

Yeast two-hybrid assays were performed as described by (Gietz and Schiestl, 2007; Lichtblau et al., 2022). C-terminal fragments of bHLH104 and b104^dPAA were cloned into the Gateway-compatible yeast vectors pACT2-GW and pGBKT7-GW. Primers used are listed in Supplementary Table 1. Full-length BTS, BTSL1, and BTSL2 constructs were described in (Lichtblau et al., 2022).

Yeast strain AH109 was co-transformed with plasmid pairs using the lithium acetate (LiAc) method and selected on synthetic dropout (SD) medium lacking leucine and tryptophan (SD–Leu–Trp). Colonies were cultured overnight in liquid SD–Leu–Trp medium, adjusted to an OD₆₀₀ of 1.0, and spotted (10 µl) in tenfold dilution series onto SD medium lacking leucine, tryptophan, and histidine (SD–Leu–Trp–His). To suppress autoactivation, SD–Leu–Trp–His plates supplemented with 15 mM 3-amino-1,2,4-triazole (3-AT) were used specifically for assays involving BTSL2. Plates were incubated at 30°C, and growth was monitored for up to 14 days.

### UbiGate Assay

Ubiquitination assays were performed using the UbiGate system as described previously (Kowarschik et al., 2018). Constructs for generating transgenic lines were assembled using the Golden Gate assembly. The following modules were assembled into pGG-pET28 destination vector: 6×His-tagged ubiquitin, E1 ligase (*Arabidopsis thaliana* UBA1), E2 ligase (Arabidopsis thaliana UBC8), E3 ligase (BTS-HC), substrate (*BHLH105* and *BHLH104* coding sequences were codon-optimized for E. coli expression using GeneArt Gene Optimizer software and synthesized by GeneArt Gene Synthesis, Thermo Fisher Scientific). Primers used are listed in Supplementary Table 1. Assembled vectors were transformed into *Escherichia coli Rosetta (DE3)* and expressed by induction with 1 mM IPTG and 3% ethanol at 28°C for 3 hours. Cells were harvested by centrifugation and stored at –80°C. Protein extraction and sample preparation was performed as previously described in (Ortmann et al., 2023).

Proteins were separated by SDS-PAGE using TGX precast gels (Bio-Rad) and transferred to 0.2 µm nitrocellulose membranes with the Trans-Blot Turbo system (Bio-Rad). Membranes were blocked with 5% (w/v) non-fat dry milk in TBST and probed with the following primary antibodies: HRP-conjugated anti-Myc (Thermo Fisher Scientific, R951-25-HRP), anti-GST (clone 8-326, Thermo Fisher Scientific) followed by HRP-conjugated secondary antibody (Jackson ImmunoResearch), HRP-conjugated Strep-Tactin (Fisher Scientific), and HRP-conjugated anti-6×His (Thermo Fisher Scientific, MA1-21315-HRP). Signals were detected using the Amersham ECL Prime kit (Cytiva) and visualized with the ChemiDoc MP Imaging System (Bio-Rad).

### Statistical Analysis and Data Visualization

All statistical analyses and data visualization were performed using RStudio (version 2025.04.1). Phenotyping data were visualized using ggplot2. Gene expression data were Z-score normalized using the scale() function in R, and gene expression heatmaps were created using the pheatmap package to visualize normalized expression levels across tissues and experimental conditions. For ICP-MS data visualization, the ggPlantMap package (Jo and Kajala, 2024) was used.

Statistical comparisons were performed using two-way ANOVA followed by Tukey’s Honest Significant Difference (HSD) test for multiple comparisons. A p-value threshold of 0.05 was considered significant.

## Acknowledgements

We thank Elke Wieneke for excellent technical assistance. The authors thank the Biocenter-MS Platform, Sabine Ambrosius and Sabine Metzger, University of Cologne, for conducting and providing support in the ICP-MS measurements. The authors thank Center for Advanced Imaging (CAi) at Heinrich Heine University for providing advanced imaging facility. DB and VV were members of the international graduate school iGRAD-Plant, Düsseldorf (the DFG International Research Training group 2466). Chat GPT 4.0 has been used to edit and shorten text, and correct and improve language.

## Author contributions

DB, PB planned and designed the research. DB, SW, MK, VLVS performed experiments and analysed data. JLE provided materials and protocol. DB wrote the manuscript. All authors read, corrected and approved the manuscript.

## Data Availability

The RNA-seq data generated in this study have been deposited in ArrayExpress at EMBL-EBI under accession number E-MTAB-16555. Confocal microscopy images have been deposited in the BioImage Archive under accession number S-BIAD2765.

---

**Supplementary Figure S1.**
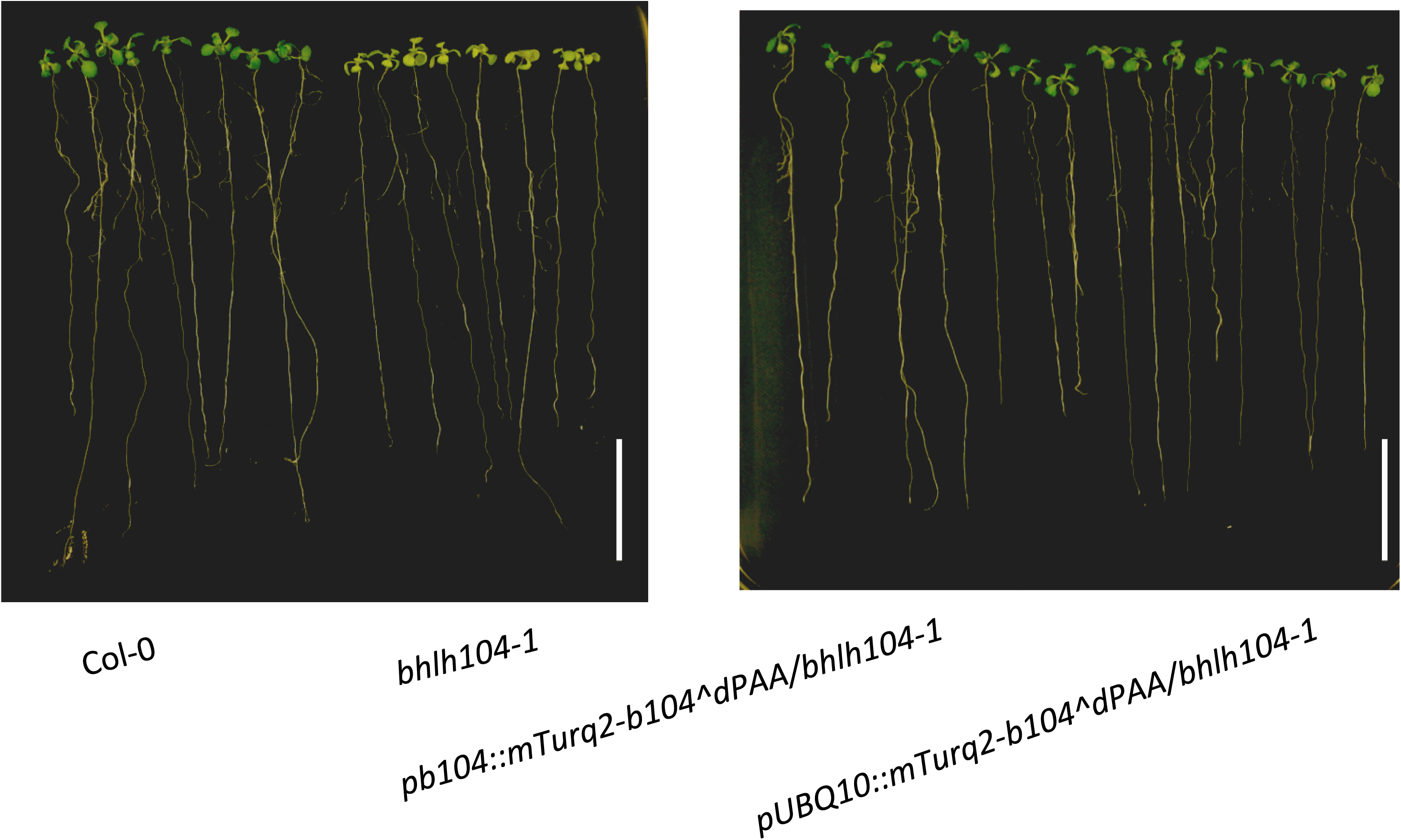
mTurquoise2-tagged bHLH104^dPAA complements the bhlh104-1 phenotype under Fe deficiency. Representative images of 12-day-old seedlings grown on iron-deficient (–Fe) medium. Left panel: wild-type Col-0 and bhlh104-1. Right panel: pb104::mTurq2-b104^dPAA/bhlh104-1 and pUBQ10::mTurq2-b104^dPAA/bhlh104-1 complementation lines. The bhlh104-1 displays chlorotic leaves compared to Col-0 under iron deficiency. Expression of mTurquoise2-tagged bHLH104^dPAA under either the native bHLH104 promoter or the UBQ10 promoter restores leaf health in the bhlh104-1 background, demonstrating that the mTurquoise2-tagged protein is biologically functional. Scale bars: 2 cm.

**Supplementary Figure S2.**
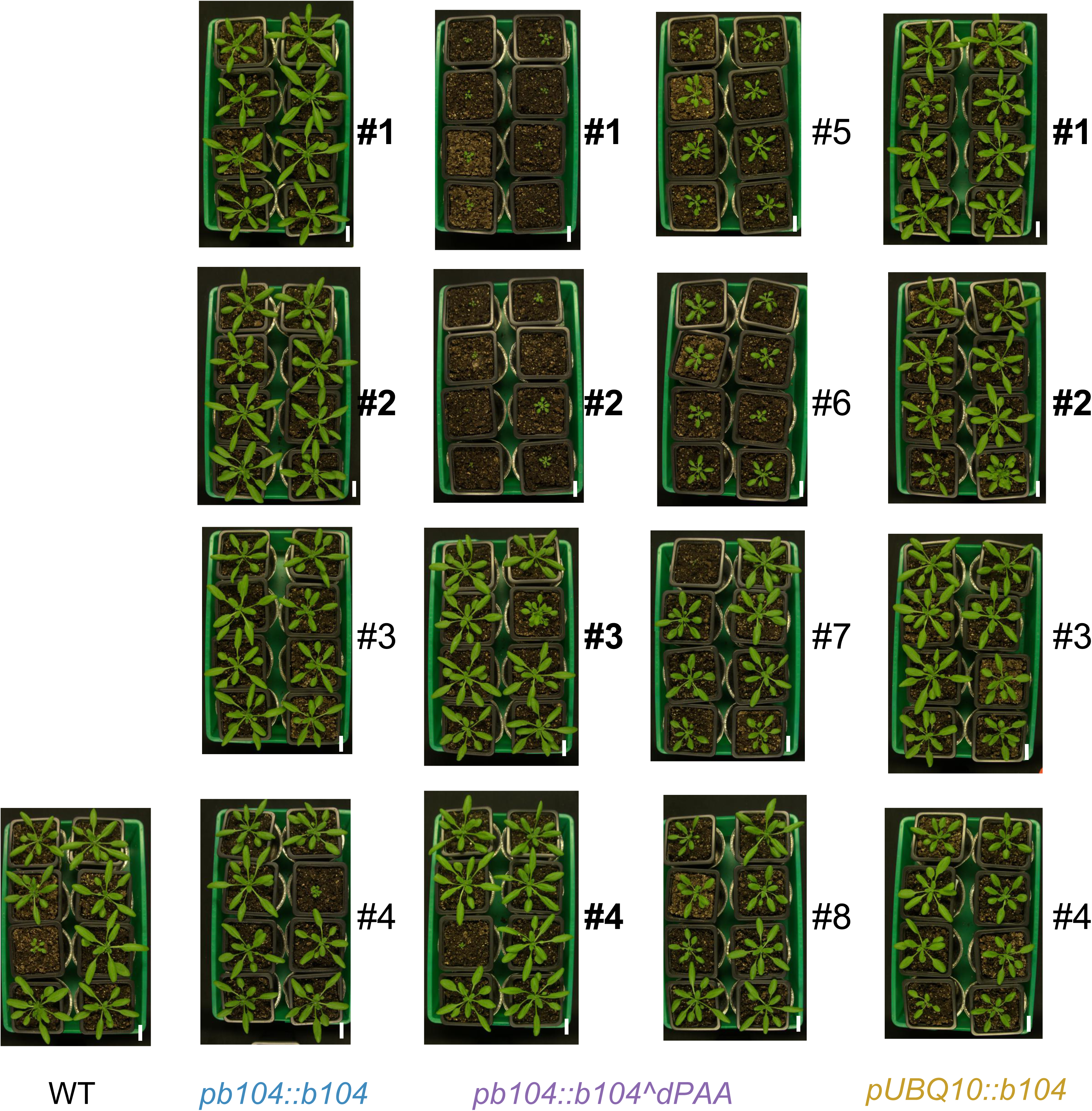
Preliminary phenotypic screen of independent transgenic Arabidopsis lines expressing bHLH104 or bHLH104^dPAA. 26 days old plants of Col-0 wild type (WT), pb104::b104 (4 independent lines), pb104::b104^dPAA (8 independent lines), and pUBQ10::b104 lines (4 independent lines), were grown under same conditions. Each tray contained eight plants derived from an independent T1 transgenic line. Lines selected for further analysis in the main study are indicated: pb104::b104 lines #1 and #2, and pb104::b104^dPAA lines #1–#4 and pUBQ10::b104 lines #1 and #2, scale bar = 2 cm.

**Supplementary Figure S3.**
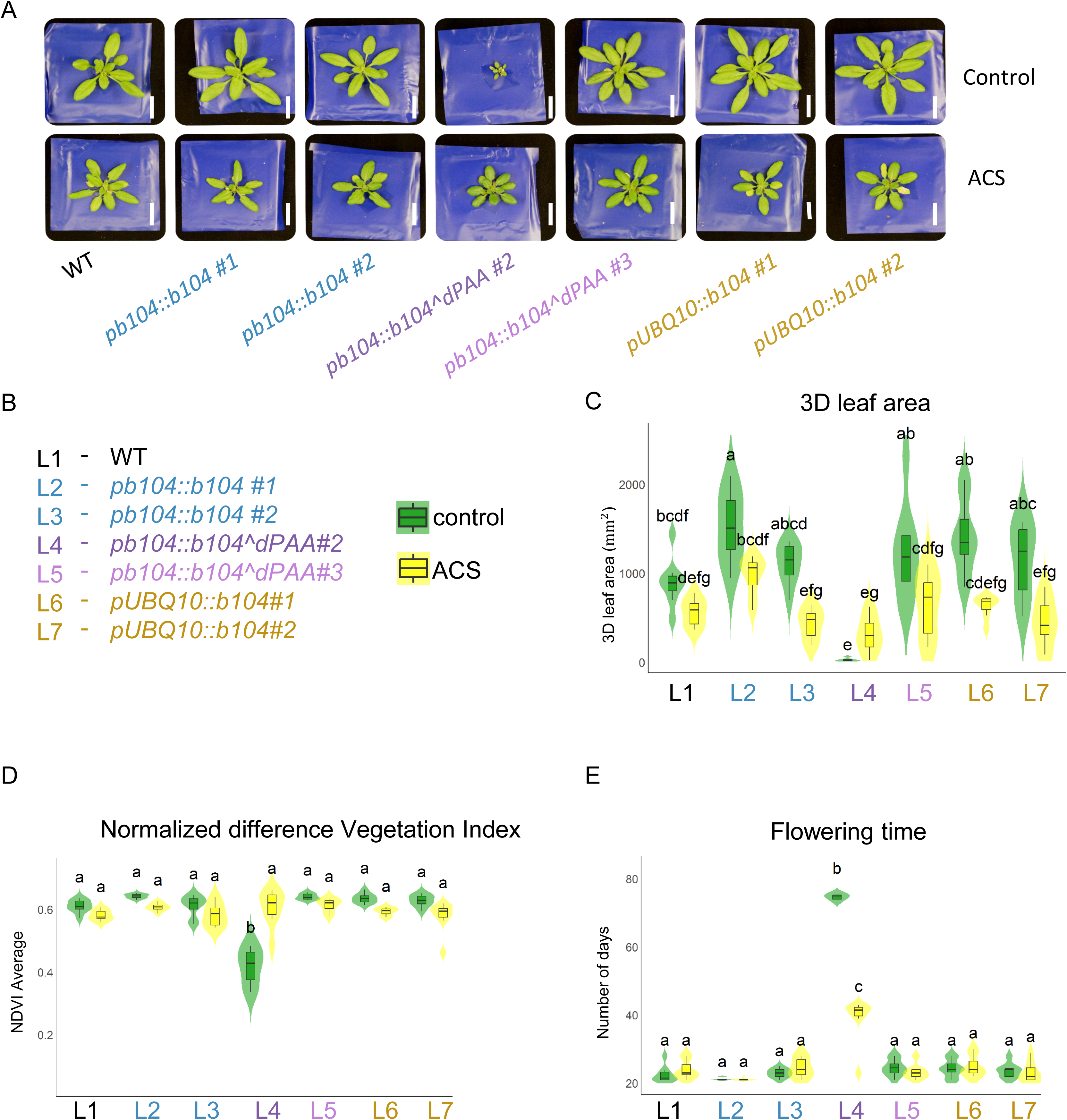
Response of bHLH104^dPAA transgenic lines to iron-limiting conditions in alkaline calcareous soil (ACS). Arabidopsis thaliana transgenic lines were grown in either control soil (pH 5.9) or alkaline calcareous soil (ACS, pH 8.1). (A) Photos of four-week-old plants grown in control and ACS soil conditions.(B) List of plant lines used in this experiment, numbered from L1 to L7. scale bar = 2 cm (C) 3D leaf area measurements: Plants were evaluated for leaf expansion and overall growth in both conditions. (D) NDVI (Normalized Difference Vegetation Index): NDVI values were measured to assess photosynthetic activity in response to ACS conditions.(E) Flowering time: Flowering time was compared between control and ACS conditions, with letters indicating statistical differences. Statistical significance was assessed by two-way ANOVA followed by Tukey’s HSD test (p < 0.05). N = 8 plants per line.

**Supplementary Figure S4.**
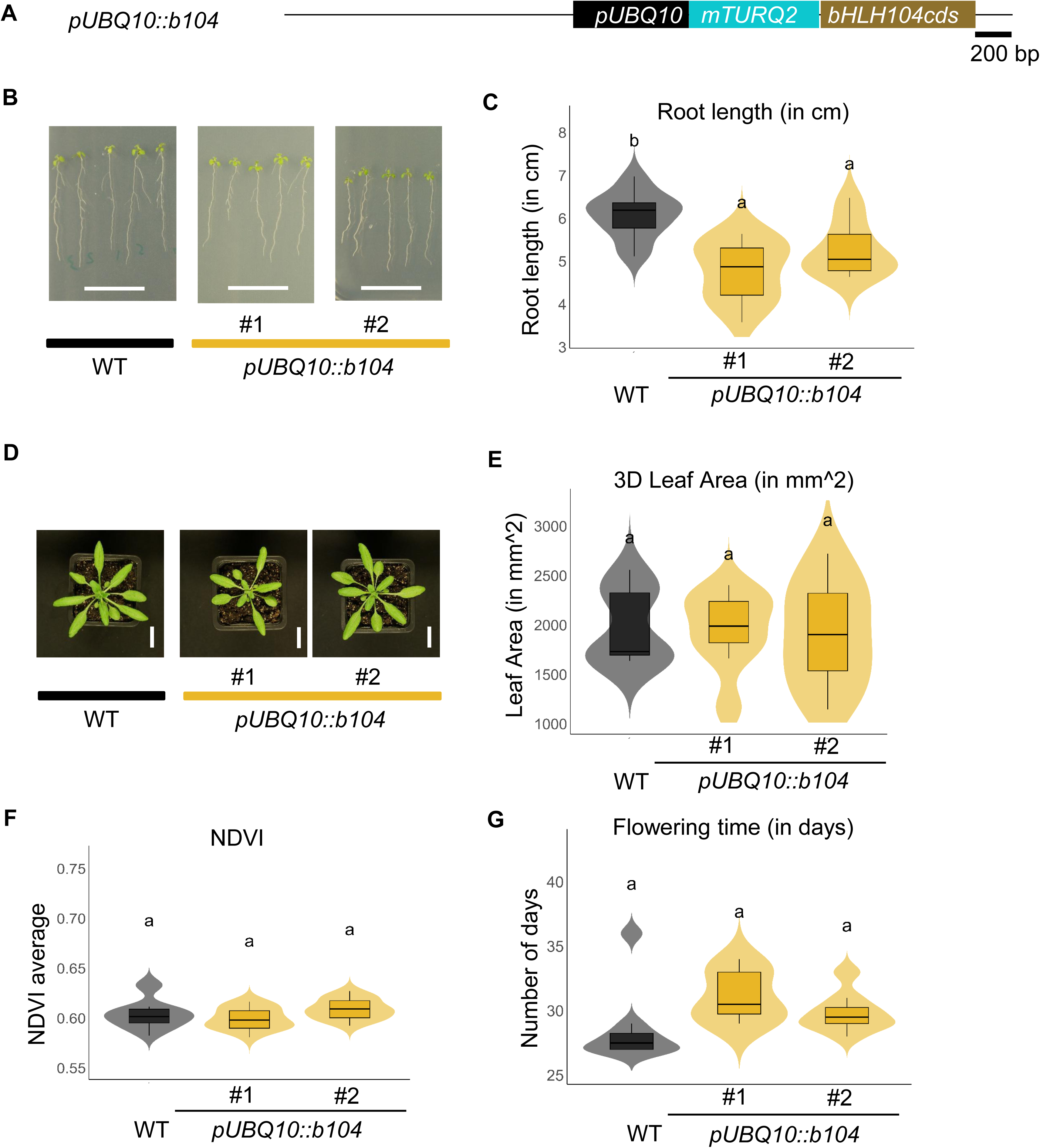
Constitutive expression of bHLH104 did not alter growth and physiological traits in Arabidopsis. (A) Schematic representation of the construct used to generate transgenic lines expressing full-length bHLH104 (pUBQ10::bHLH104). The construct is driven by the ubiquitin 10 (UBQ10) promoter and is fused at the N-terminus to the mTurquoise2 fluorescent tag. Scale bar = 200 bp. (B-C) Photos and root length quantification data of 12-day-old seedlings grown vertically on Hoagland medium supplemented with iron (+Fe). Scale bar =2 cm (D) Representative images of 4-week-old Arabidopsis plants grown in soil. Shown are Col-0 (WT) and two independent pUBQ10::b104 lines (#1, #2). Scale bar = 2 cm (E–F) Quantitative analysis of aboveground phenotypes acquired using the Phenospex PlantEye system: (E) 3D leaf area (mm²); (F) NDVI (Normalized Difference Vegetation Index). (G) Flowering time, measured manually as the number of days to bolting. Data represent means ± SD (n = 7–12 biological replicates per line). Statistical analysis was performed using two-way ANOVA followed by Tukey’s post hoc test. Different letters indicate statistically significant differences at p < 0.05.

**Supplementary Figure S5.**
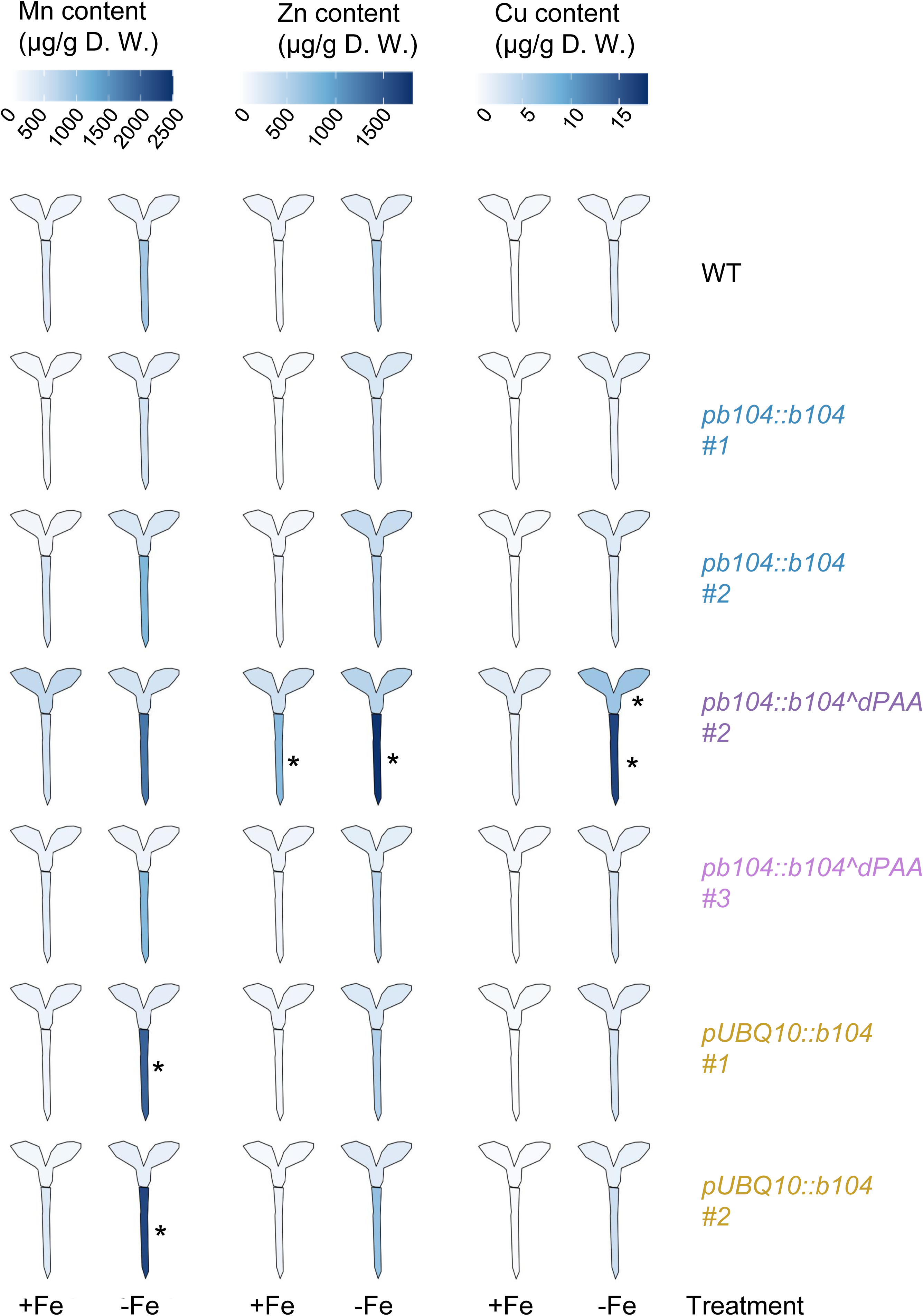
Elemental content analysis of Mn, Zn, and Cu in 10-day old Arabidopsis seedlings. Heatmap showing the distribution of Mn, Zn, and Cu content (in µg/g dry weight) across different Arabidopsis seedlings, including wild-type (WT), pb104::b104 (lines #1, #2), pb104::b104^dPAA (lines #2, #3), pUBQ10::b104 (lines #1, #2). The color scale indicates the range of elemental concentrations from low (light blue) to high (dark blue). Seedlings were grown in Hoagland media with or without 50 mM Fe supplementation. Data were collected from 10-day-old seedlings, with elemental content measurements obtained via ICP-MS analysis. Three biological replicates per line were used. The plot was generated using the ggPlantMap package in RStudio. Asterisks indicate genotypes that differ significantly from WT within the same tissue and treatment condition (p < 0.05), as determined by two-way ANOVA and Tukey’s post hoc test (n = 3 for each group).

**Supplementary Figure S6.**
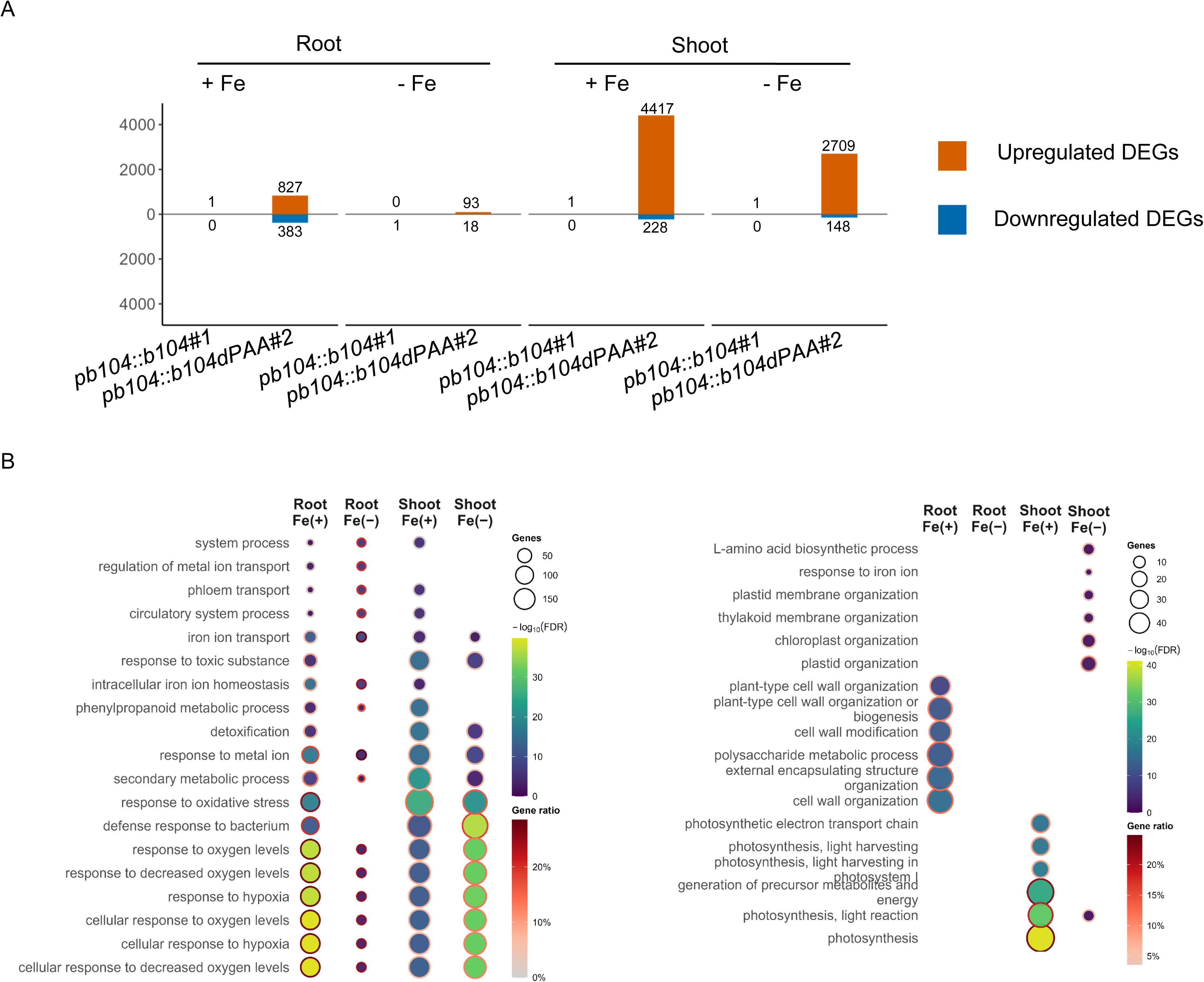
Differential expression and GO enrichment across seedling tissues. (A) Counts of differentially expressed genes (DEGs) for each line vs WT in root and shoot under Fe(+) and Fe(−). Orange = up-regulated; blue = down-regulated; numbers above/below bars indicate counts for each category. Total DEGs for each condition are the sum of up-regulated and down-regulated counts. DEGs were defined at FDR ≤ 0.05 and |log2FC| ≥ 0.58. X-axis labels show the abbreviated line IDs. (B) GO Biological Process enrichment for the up-regulated (left panel) and down-regulated (right panel) DEG sets. Columns correspond to tissue/Fe combinations (Root Fe(+), Root Fe(−), Shoot Fe(+), Shoot Fe(−)). Each dot represents a significant term (BH FDR ≤ 0.05); dot size = number of genes in the term, fill = −log10(FDR), and ring color = GeneRatio. Terms plotted are the top-ranked per panel; the gene universe was all genes tested in the corresponding comparison.

**Supplementary Figure S7.**
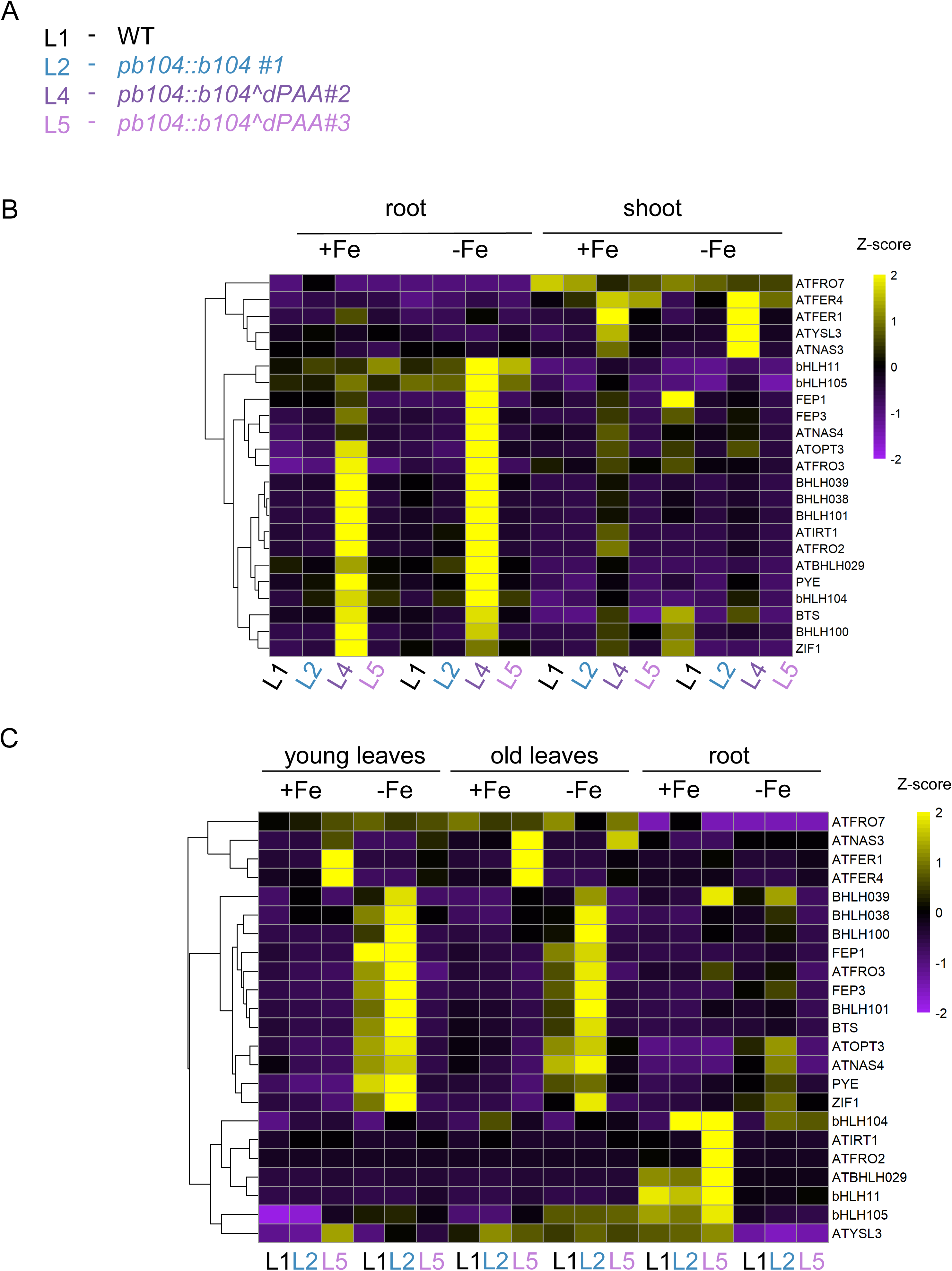
Expression profiles of 23 selected Fe homeostasis genes involved in Fe acquisition and mobilization. (A) Plant lines used In this experiment (B) Clustered heatmap showing expression profiles of 23 Fe homeostasis genes (gene selection based on Ngigi et al., 2025; Schwarz et al., 2020) obtained using RNA sequencing in root and shoot tissues under +Fe and –Fe conditions. Arabidopsis seedlings were grown for 10 days after sowing (DAS) under either iron-sufficient (+Fe) or iron-deficient (–Fe) conditions. (C) Clustered heatmap showing expression profiles of 23 Fe homeostasis genes (gene selection based on Ngigi et al., 2025; Schwarz et al., 2020) obtained using RNA sequencing in young leaves, old leaves, and roots under both +Fe and –Fe conditions. Arabidopsis plants were grown hydroponically under iron-sufficient (+Fe) conditions for 27 days, followed by a 3-day treatment with either +Fe or –Fe media. Heatmap was generated in RStudio version 4.4.1 using calculated Z-scores of normalized gene expression values. Purple indicates low expression and yellow indicates high expression.

**Supplementary Figure S8.**
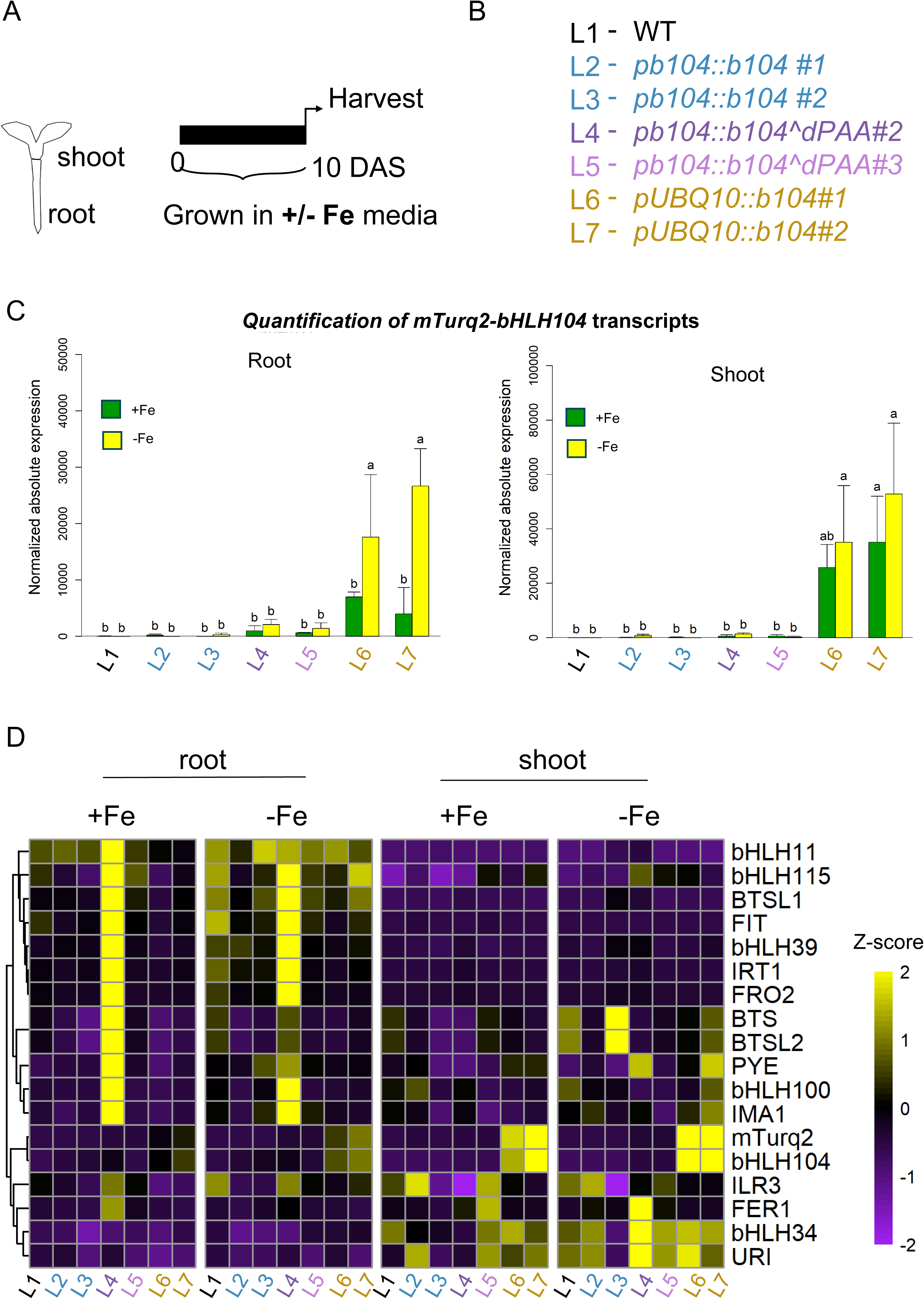
Targeted gene expression profiling in seedlings. (A) Schematic representation of the experimental design. Arabidopsis seedlings were grown for 10 days after sowing (DAS) under either iron-sufficient (+Fe) or iron-deficient (–Fe) conditions. Shoot and root tissues were collected separately for RT-qPCR analyses. (B) Genotypes used in this study. Abbreviations of lines used: L1, wild-type Col-0; L2, pb104::mTurquoise2-bHLH104 line #1; L3, pb104::mTurquoise2-bHLH104 line #2; L4, pb104::mTurquoise2-b104^dPAA line #2; L5, pb104::mTurquoise2-b104^dPAA line #3; L6, pUBQ10::mTurquoise2-bHLH104 line #1; L7, pUBQ10::mTurquoise2-bHLH104 line #2. (C) Quantification of mTurq2-tagged bHLH104 transgenes in indicated plant lines using RT-qPCR Letters indicating statistical differences. Statistical significance was assessed by two-way ANOVA followed by Tukey’s HSD test (p < 0.05). N = 3 per group. (D) Clustered heatmaps showing the expression profiles of selected Fe homeostasis genes obtained using RTqPCR. Heatmaps were generated in RStudio version 4.4.1 using calculated Z-scores of normalized gene expression values across root and shoot tissues for both +Fe and –Fe conditions. Purple indicates low expression and yellow indicates high expression.

**Supplementary Figure S9.**
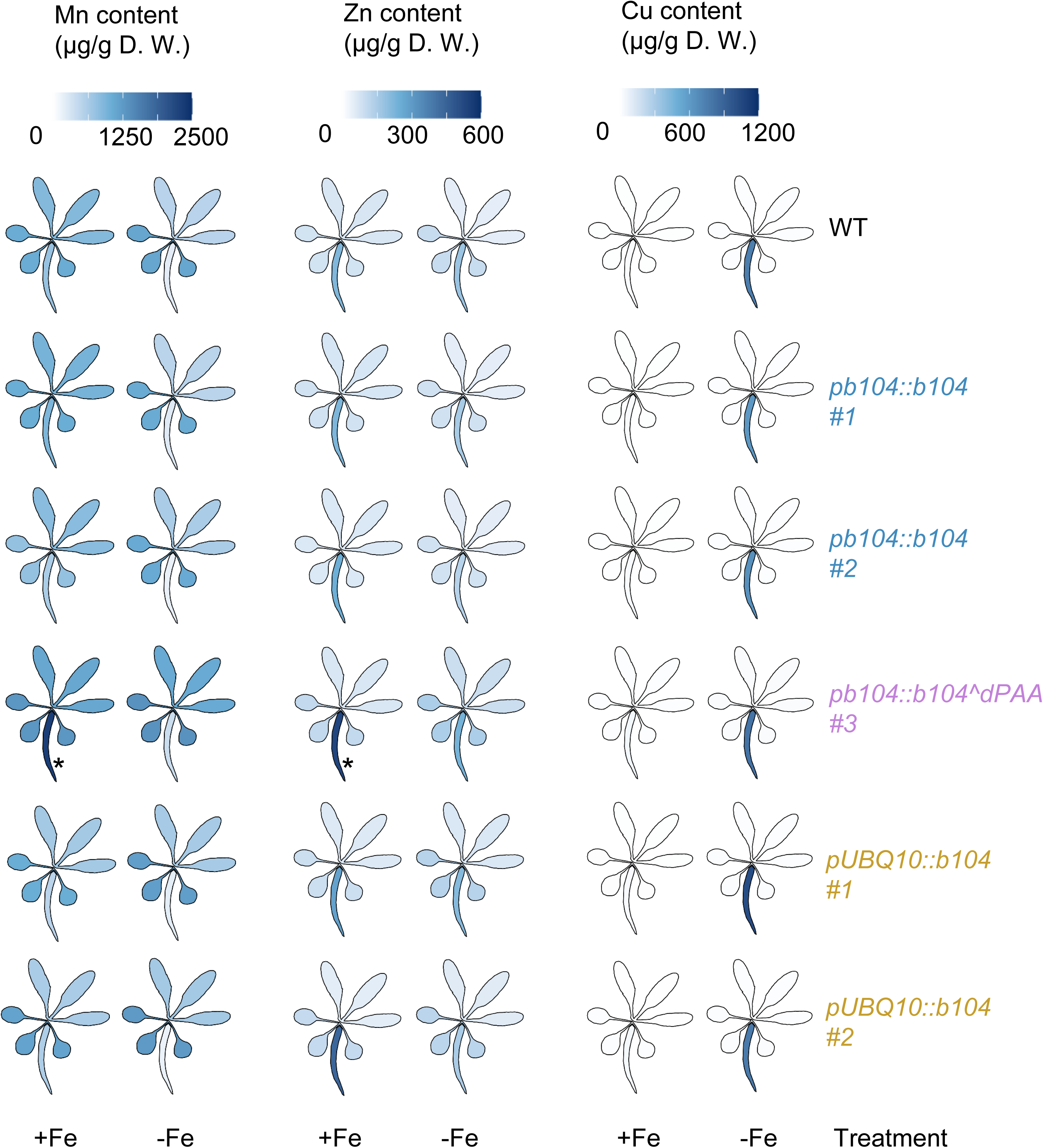
Elemental content analysis of Mn, Zn, and Cu in mature Arabidopsis thaliana. Heatmap showing the distribution of iron (Fe) content (in µg/g dry weight) in young leaves, old leaves, and roots of Arabidopsis transgenic lines, including wild-type (WT), pb104::b104 (lines #1, #2), pb104::b104^dPAA (line #3), pUBQ10::b104 (lines #1, #2). Plants were grown hydroponically under iron-sufficient (+Fe) conditions for 27 days, followed by a 3-day treatment with either +Fe or –Fe media. The color scale indicates the iron content, with darker blue shades representing higher iron concentrations. Data represent the mean values from three biological replicates (n = 3). Heatmap intensity reflects iron concentration (µg/g dry weight) measured by ICP-MS. The plot was generated using the ggPlantMap package in RStudio. Asterisks indicate genotypes that differ significantly from WT within the same tissue and treatment condition (p < 0.05), as determined by two-way ANOVA and Tukey’s post hoc test (n = 3 for each group).

**Supplementary Figure S10.**
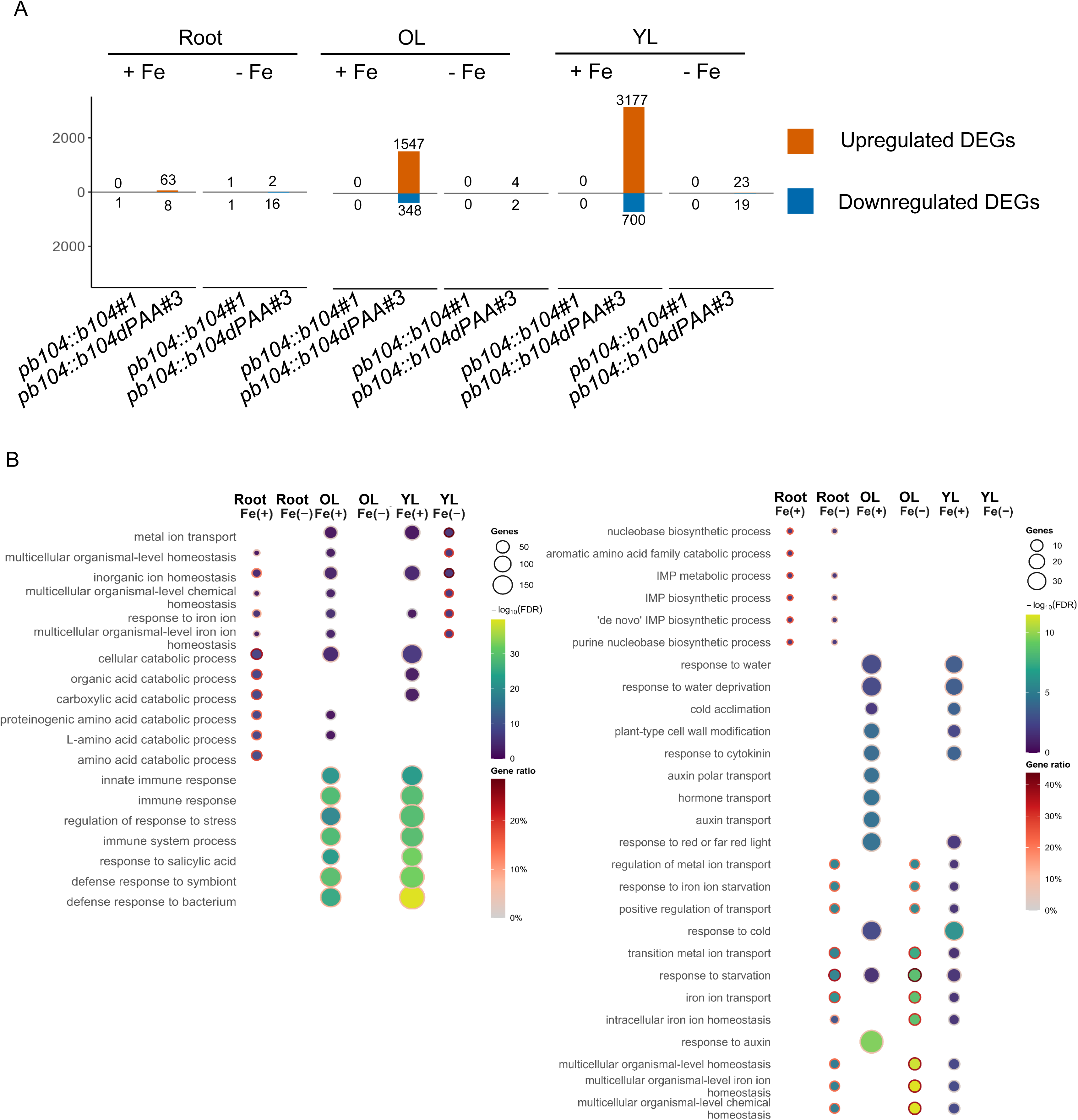
Differential expression and GO enrichment across reproductive stage tissues. (A) Numbers of differentially expressed genes (DEGs) for each line vs WT in root (Root), young leaves (YL), and old leaves (OL) under Fe(+) and Fe(−). Orange bars show up-regulated DEGs; blue bars show down-regulated DEGs; values above/below bars are counts for each category. Total DEGs for each condition are the sum of up- regulated and down-regulated counts. DEGs were defined at FDR ≤ 0.05 and |log2FC| ≥ 0.58. (B) GO Biological Process enrichment for the up-regulated (left panel) and down-regulated (right panel) DEG sets. Columns correspond to tissue/Fe combinations (Root, YL, OL × Fe (+), Fe (−)). Each dot represents a significant term (BH FDR ≤ 0.05); dot size = number of genes in the term, fill = −log10(FDR), and ring color = Gene Ratio. Terms shown are the top-ranked per panel; the gene universe comprised all genes tested in the corresponding comparison. Abbreviations: YL, young leaves; OL, old leaves.

**Supplementary Figure S11.**
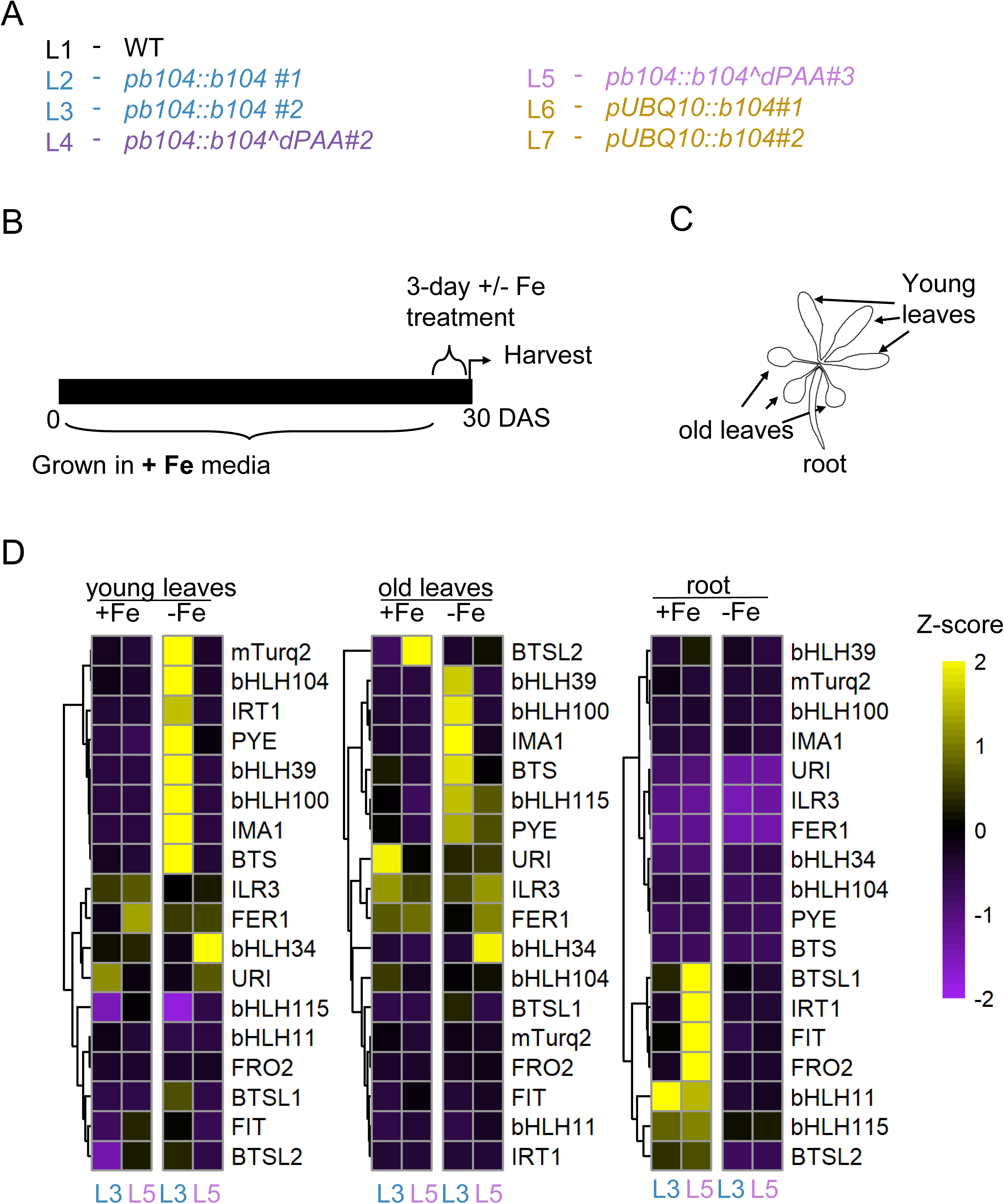
Targeted gene expression profiling in reproductive-stage plants. (A) Plant lines used In this experiment. Abbreviations of lines used: L1, wild-type Col-0; L2, pb104::mTurquoise2-bHLH104 line #1; L3, pb104::mTurquoise2-bHLH104 line #2; L4, pb104::mTurquoise2-b104^dPAA line #2; L5, pb104::mTurquoise2-b104^dPAA line #3; L6, pUBQ10::mTurquoise2-bHLH104 line #1; L7, pUBQ10::mTurquoise2-bHLH104 line #2. (B) Experimental design for the dataset shown. Arabidopsis plants were grown hydroponically under iron-sufficient (+Fe) conditions for 27 days, followed by a 3-day treatment with either +Fe or –Fe media. (C) Schematic representation of tissue separation used for RT-qPCR analysis. Young and old leaves were separated from the roots to examine tissue-specific iron accumulation and gene expression. (D) Clustered heatmaps showing the expression profiles of selected Fe homeostasis genes conditions obtained using RTqPCR in young leaves, old leaves, and roots under both +Fe and –Fe. Data represent the mean of three biological replicates (n = 3). Z-scores were calculated from normalized RT-qPCR expression values and visualized in RStudio version 4.4.1. Yellow indicates high expression; purple indicates low expression.

**Supplementary figure S12.**
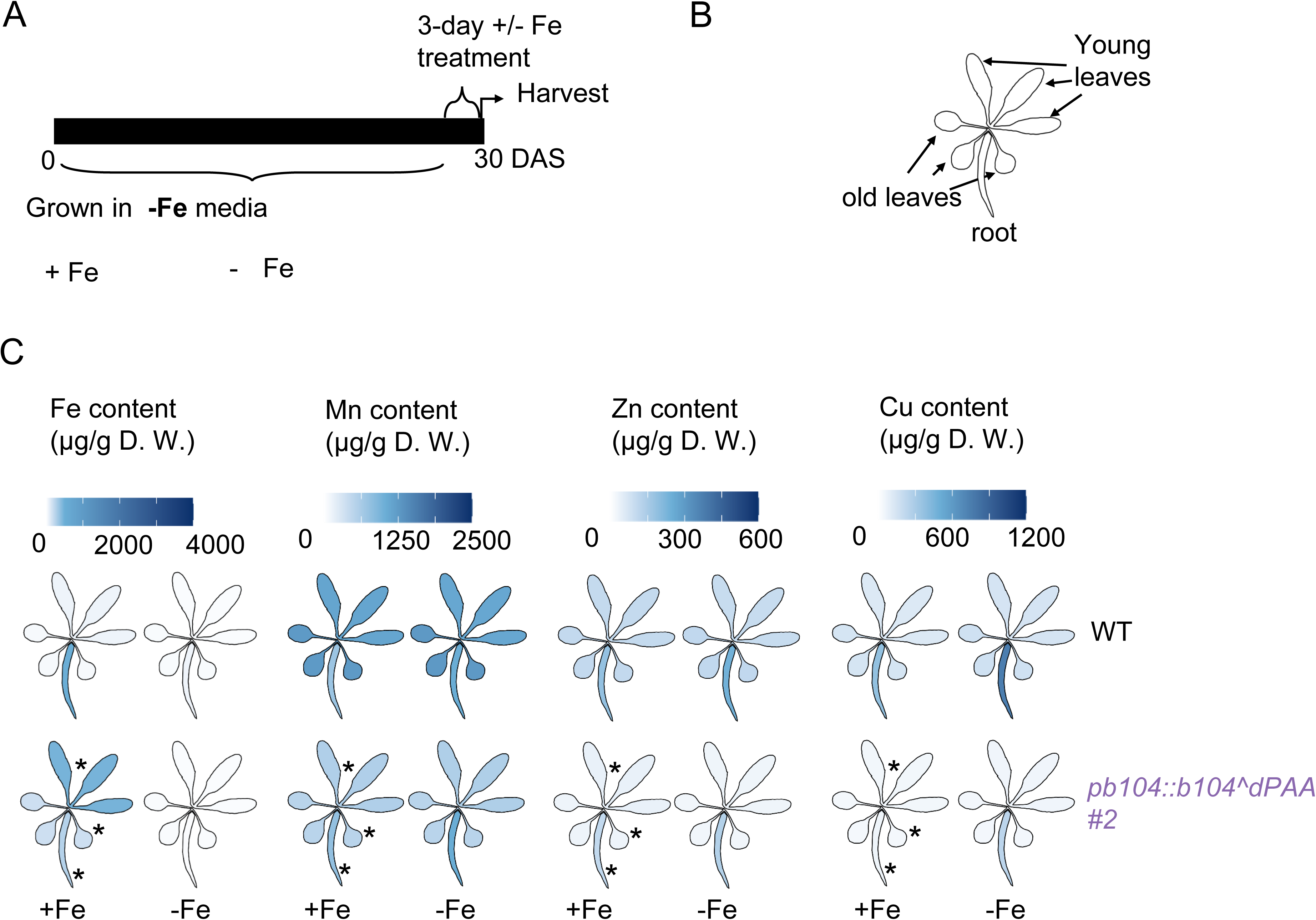
Tissue-specific metal accumulation in dPAA line #2 grown under iron-deficient conditions. (A) Experimental setup: Col-0 and pb104::b104^dPAA line #2 seedlings were grown hydroponically in Fe-deficient (–Fe) media for 27 days, followed by a 3-day treatment with either +Fe or –Fe media. Tissue collection occurred at 30 days after sowing (DAS). (B) Tissue layout: Young leaves, old leaves, and roots were separated to examine tissue-specific metal accumulation (Fe, Mn, Zn, Cu).(C) Heatmap of metal content: The distribution of Fe, Mn, Zn, and Cu content (in µg/g dry weight) in dPAA line #2 and Col-0. The color scale reflects metal concentrations, with darker blue shades indicating higher content. pb104::b104^dPAA line #2 exhibited significant Fe accumulation in young leaves after Fe resupply, in contrast to Col-0, where most Fe remained in the roots. Data represent the mean values from three biological replicates (n = 3). Elemental content was measured by ICP-MS. The plot was generated using the ggPlantMap package in RStudio. Asterisks indicate genotypes that differ significantly from WT within the same tissue and treatment condition (p < 0.05), as determined by two-way ANOVA and Tukey’s post hoc test (n = 3 for each group).

**Supplementary Figure S13.**
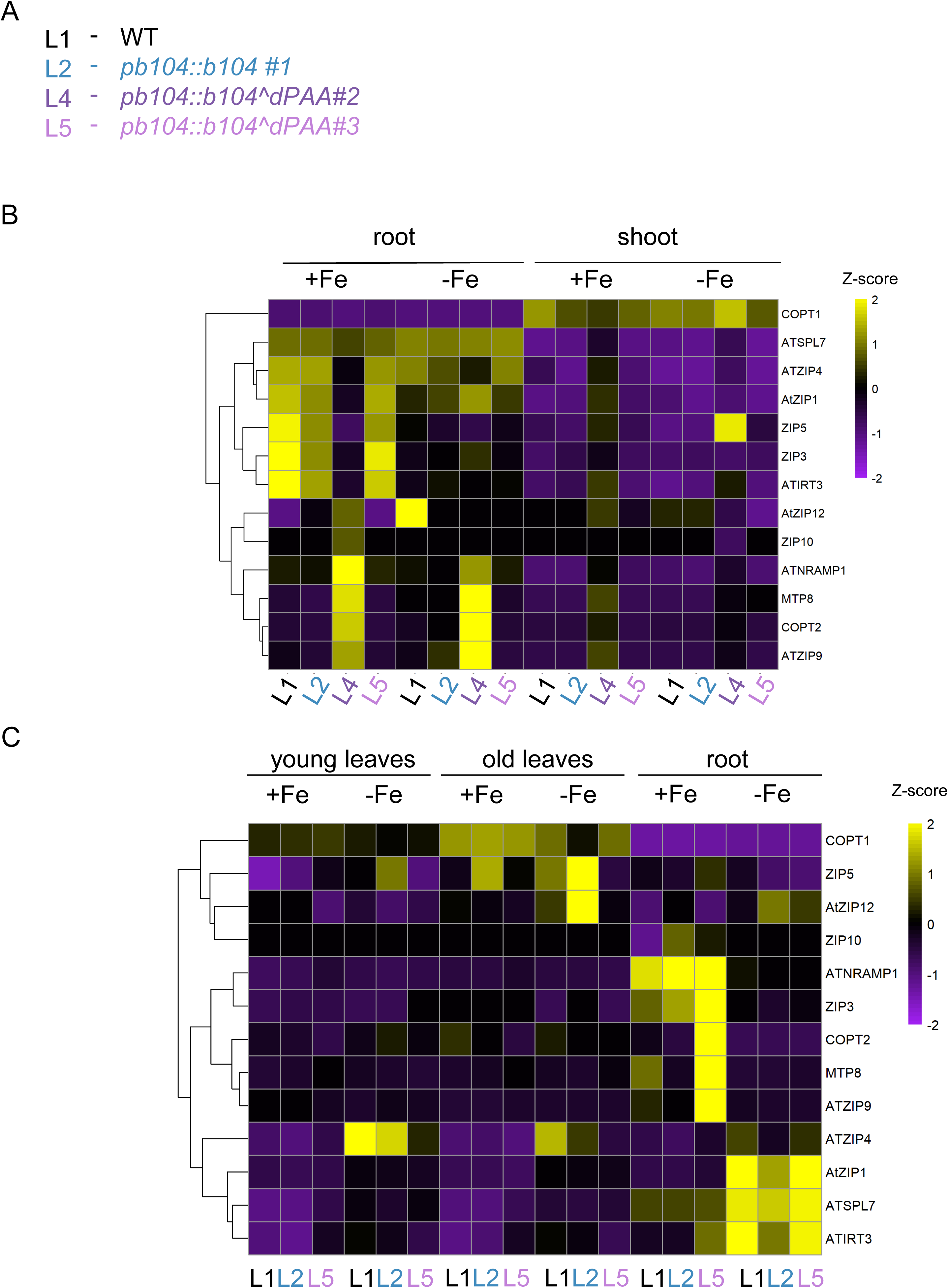
Expression profiles of genes involved in Mn, Zn, and Cu homeostasis. (A) Plant lines used In this experiment (B) Clustered heatmap showing expression profiles from RNA sequencing experiment of genes involved in manganese (Mn), zinc (Zn), and copper (Cu) homeostasis across root and shoot tissues under +Fe and –Fe conditions. Arabidopsis seedlings were grown for 10 days after sowing (DAS) under either iron-sufficient (+Fe) or iron-deficient (–Fe) conditions. (C) Clustered heatmap showing expression profiles from RNA sequencing experiment of genes involved in manganese (Mn), zinc (Zn), and copper (Cu) homeostasis in young leaves, old leaves, and roots under both +Fe and –Fe conditions. Arabidopsis plants were grown hydroponically under iron-sufficient (+Fe) conditions for 27 days, followed by a 3-day treatment with either +Fe or –Fe media. Genes include metal transporters and the Cu homeostasis regulator SPL7. Heatmap was generated in RStudio version 4.4.1 using calculated Z-scores of normalized gene expression values. Purple indicates low expression and yellow indicates high expression.

**Supplementary Figure S14.**
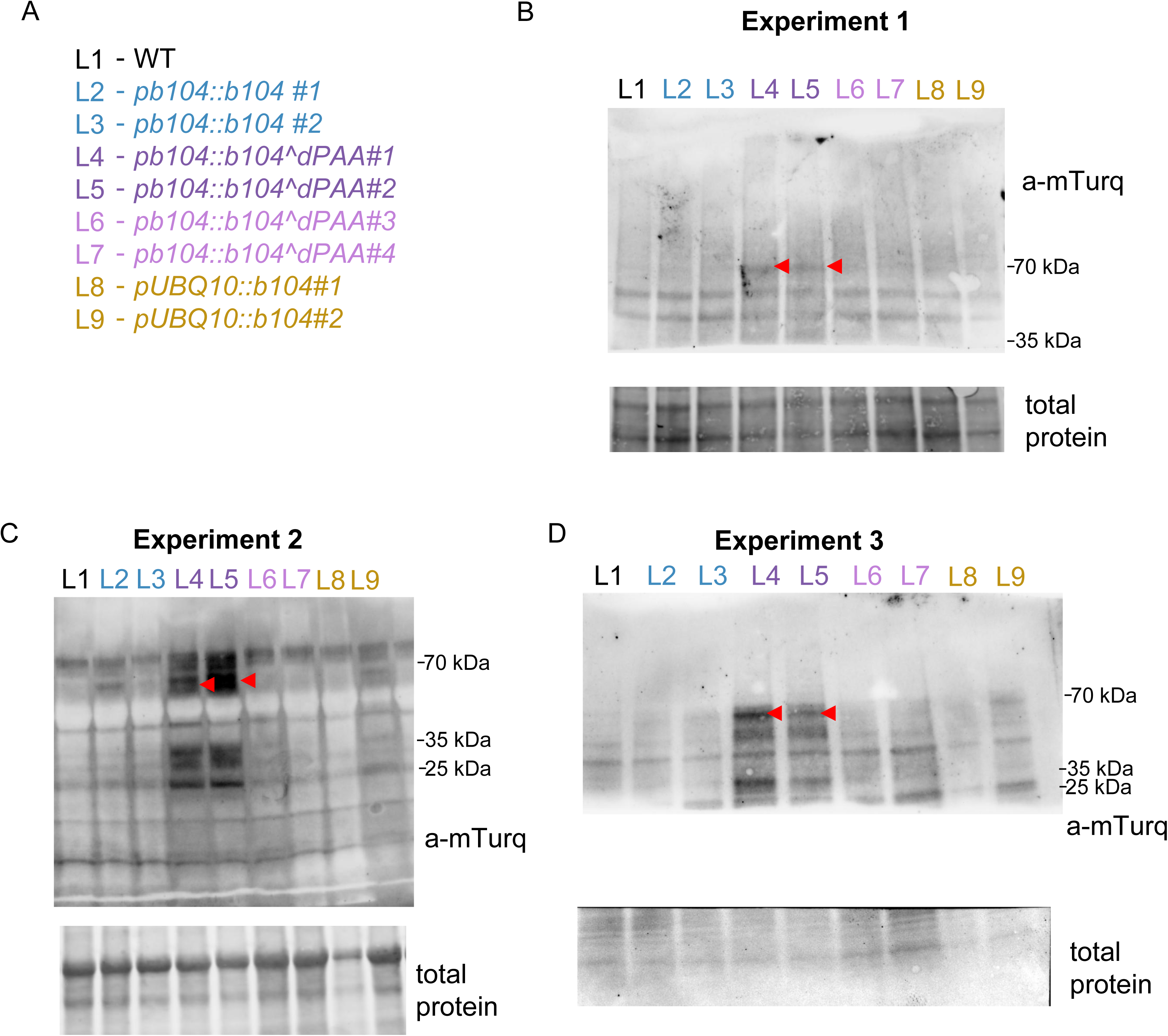
Additional independent replicates and anti-mTurq immunoblots for detection of mTurquoise2-bHLH104 and -bHLH104^dPAA protein in various transgenic lines. Genotypes and transgenic lines used in this experiment. Genotypes and transgenic lines used in this experiment. The mTurquoise2 fluorescent tag is fused at the N-terminus of bHLH104 and b104^dPAA in all transgenic lines. Experiment 1: immunoblot of total protein extracted from roots of 10-day-old seedlings (see also main text figure, Fig. 5B). Experiment 2: immunoblot of total protein extracted from 10-day-old whole seedlings. Experiment 3: immunoblot of total protein extracted from 6-day-old whole seedlings. For each experiment, top panels show the anti-mTurquoise2 blot and bottom panels show total protein stain as loading control. Molecular weight markers (kDa) are indicated on the right of each blot. Red arrowheads indicate the expected band position for the full-length mTurquoise2-bHLH104 fusion protein (∼70 kDa). Lower molecular weight bands detected by the anti-mTurquoise2 antibody likely represent truncated or partially degraded forms of the mTurquoise2-tagged protein. Across all three independent experiments, mTurquoise2-b104^dPAA protein is consistently more abundant than mTurquoise2-bHLH104, supporting that deletion of the PAA sequence stabilizes bHLH104 protein. The three experiments represent independent biological samples differing in seedling age (6 or 10 days) and tissue type (whole seedlings or roots), demonstrating the robustness of the observed protein accumulation pattern.

**Supplementary Figure S15.**
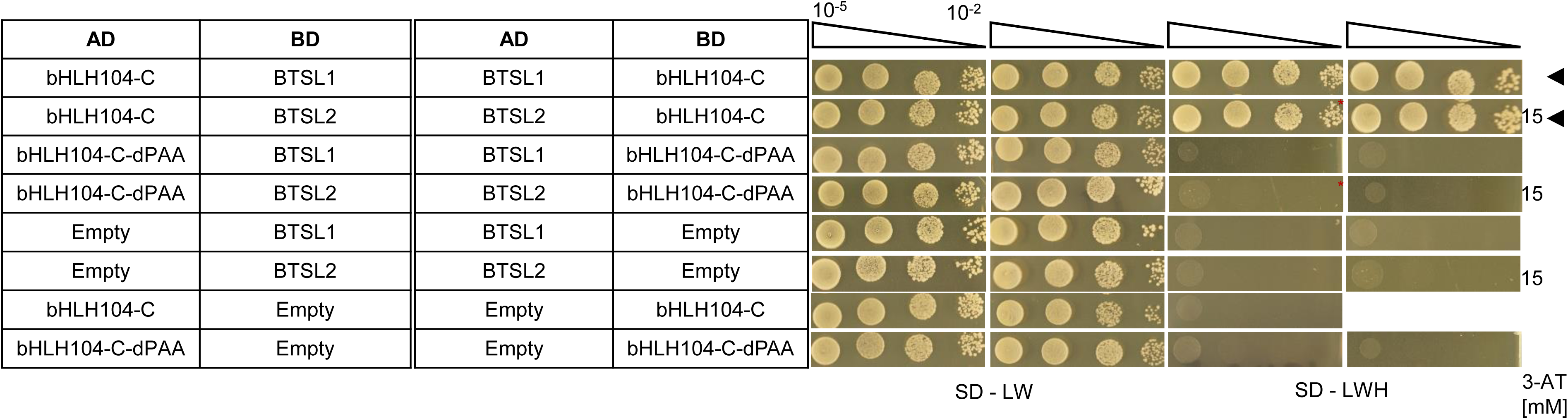
Yeast two-hybrid assay results showing protein–protein interactions between bHLH104 BTSL1 and BTSL2. bHLH104 interacts with BTSL1 and BTSL2, but not bHLH104dPAA. bHLH104^ΔPAA retains the ability to heterodimerize with full-length bHLH104, serving as a control for the presence and expression of the PAA-deleted variant. Full length bHLH104 shows autoactivation in Y2H. Hence bHLH104-C and bHLH104-C-dPAA were used. Yeast co-transformed with the AD and BD combinations were spotted in 10-fold dilution series (A600 = 10–2–10–5) on SD-LW (transformation control) and SD-LWH plates supplemented with different 3AT concentrations (conc.) as indicated (selection for protein interaction). Negative controls: empty vectors. Black arrows indicate interaction.

**Supplementary Figure S16.**
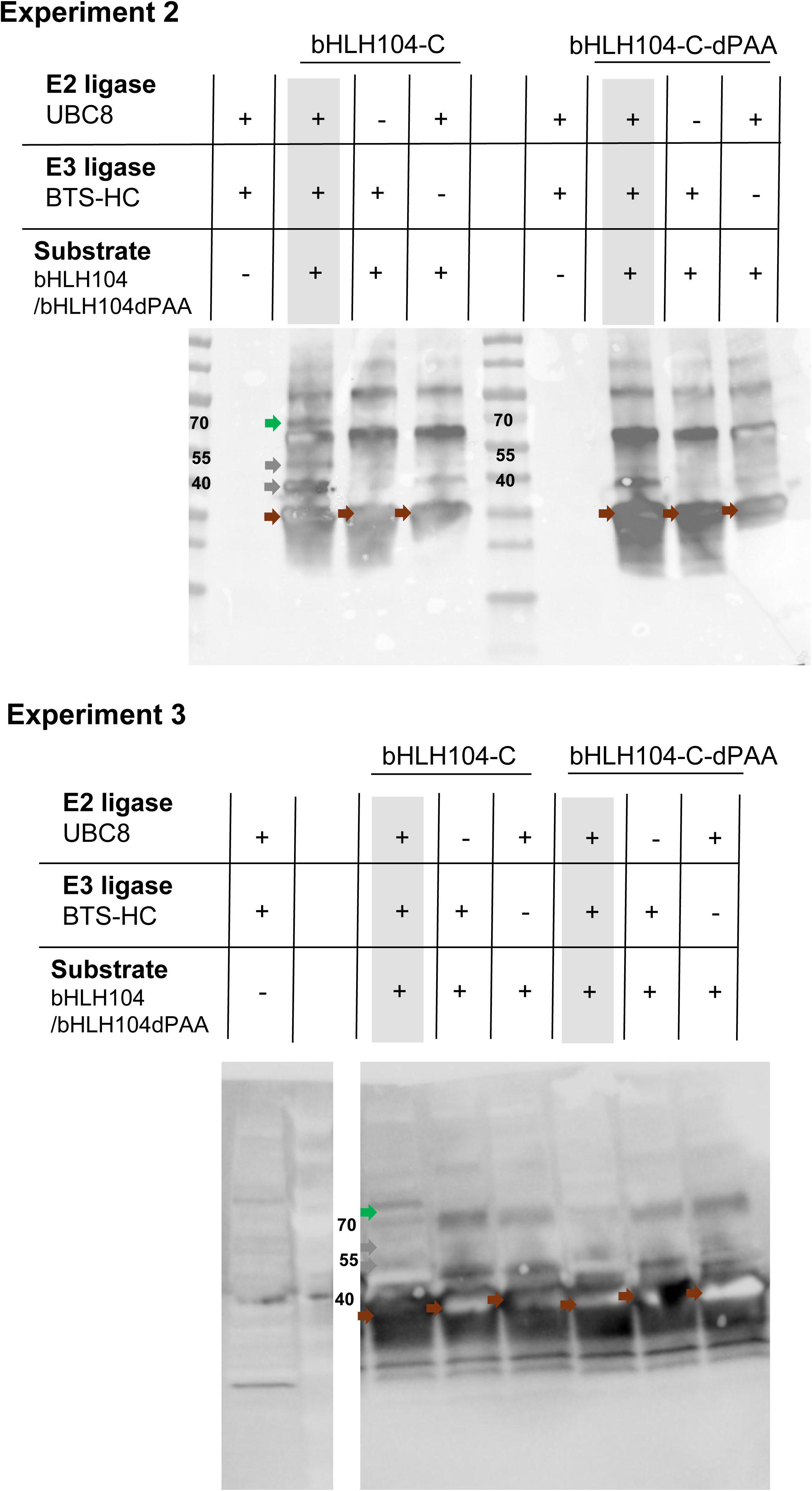
Independent replicates of the UbiGate ubiquitination assay for bHLH104. Anti-Strep immunoblots from a total of three independent UbiGate experiments (Experiment 1, see also Fig. 7B; additional Experiment 2 and Experiment 3) supporting the possible PAA-dependent ubiquitination of bHLH104 by BTS. The anti-Strep antibody detects Strep-tagged substrate protein (∼40 kDa). Brown arrows indicate non-ubiquitinated substrate. Green arrows indicate a distinct and unique higher molecular weight band (∼70 kDa), present only in the bHLH104-C sample in the presence of all components, but in none of the control samples nor the bHLH104-C^dPAA samples consistent with polyubiquitinated bHLH104-C, representing a molecular weight shift of approximately 30 kDa corresponding to the addition of three to four ubiquitin moieties. Grey arrows indicate additional higher molecular weight bands that are not considered specific, as similar bands are present across control lanes. In both replicates, the ∼70 kDa polyubiquitinated band is seen, supporting the resulting outcome shown in Figure 7B.

**Supplementary Figure S17.**
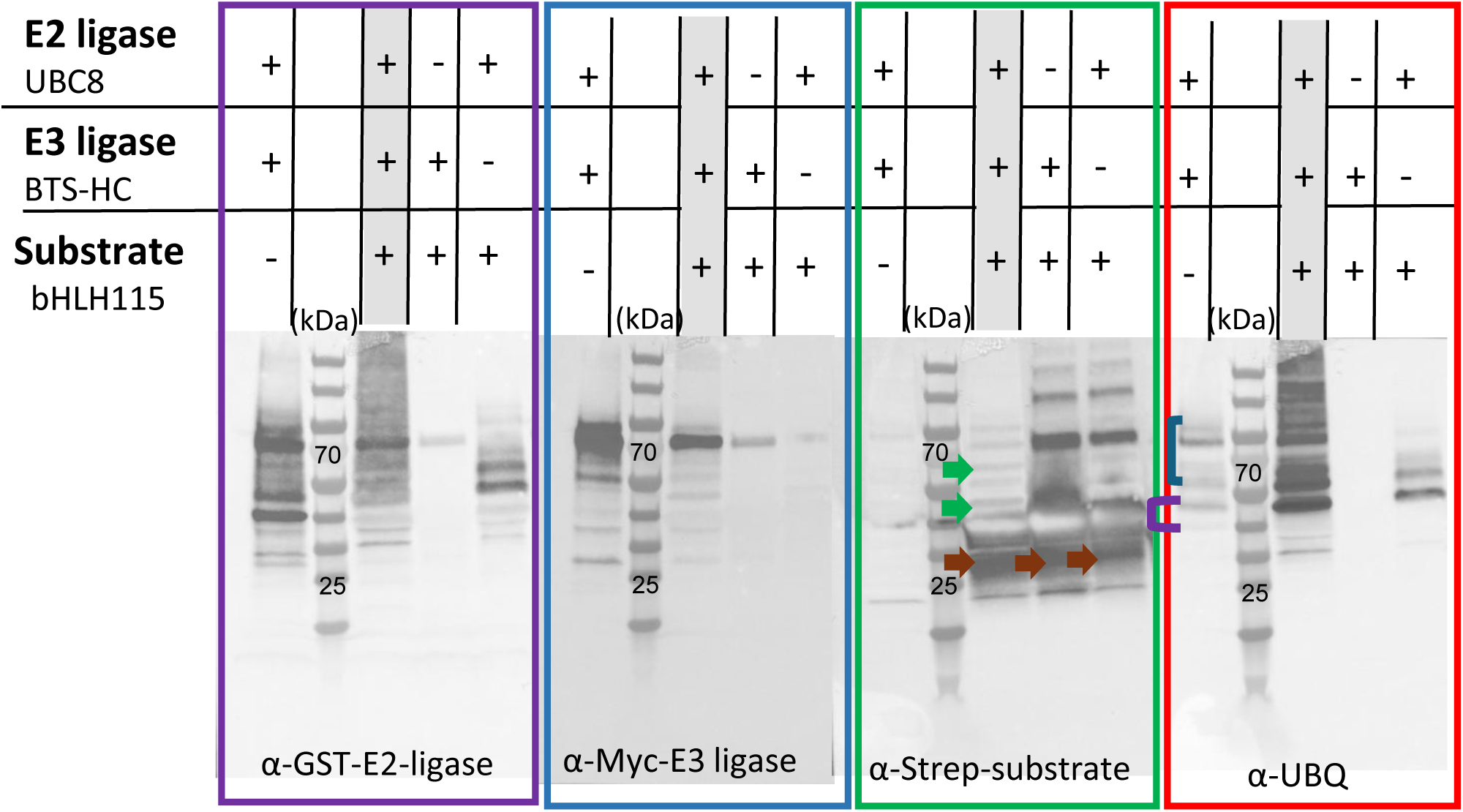
UbiGate assay showing ubiquitination of bHLH115 by BTS-HC. UbiGate ubiquitination assay testing whether bHLH115 is ubiquitinated by BTS-HC in E. coli cells. All four panels are from the same experiment, probed sequentially with different antibodies after stripping: anti-GST (E2 ligase UBC8 detection, purple box), anti-Myc (E3 ligase BTS-HC detection, blue box), anti-Strep (substrate bHLH115 detection, green box), and anti-His-UBQ (ubiquitin detection, red box). Lane compositions are indicated above: +/− denotes presence or absence of E2 ligase (UBC8), E3 ligase (BTS-HC), and substrate (Strep-tagged bHLH115). The first lane in each panel contains E2 and E3 without substrate, serving as a substrate-negative control. The anti-Strep blot (green box) shows the key substrate-specific result: brown arrows indicate the unmodified bHLH115 substrate, and green arrows indicate higher molecular weight bands present in the complete reaction lane (+E2, +E3, +substrate) that are consistent with polyubiquitinated bHLH115 forms. Several non-specific higher molecular weight bands are present across multiple lanes including controls. However, the complete reaction lane shows a distinct shift in banding pattern with clear substrate laddering compared to the E2-minus and E3-minus controls, indicating that polyubiquitination requires both E2 and E3 components. In the anti-UBQ blot (red box), the presence of all components (+E2, +E3, +substrate) produces a distinct ubiquitin banding pattern compared to controls, further supporting substrate ubiquitination. Violet bracket indicates ubiquitinated E2 ligase (UBC8). Blue bracket indicates autoubiquitinated E3 ligase (BTS-HC). These results are consistent with previous reports of BTS-dependent degradation of bHLH115 (Selote et al., 2015; Xing et al., 2021) and demonstrate the utility of the UbiGate system for testing ubiquitination of bHLH IVc proteins.

**Supplementary Figure S18.**
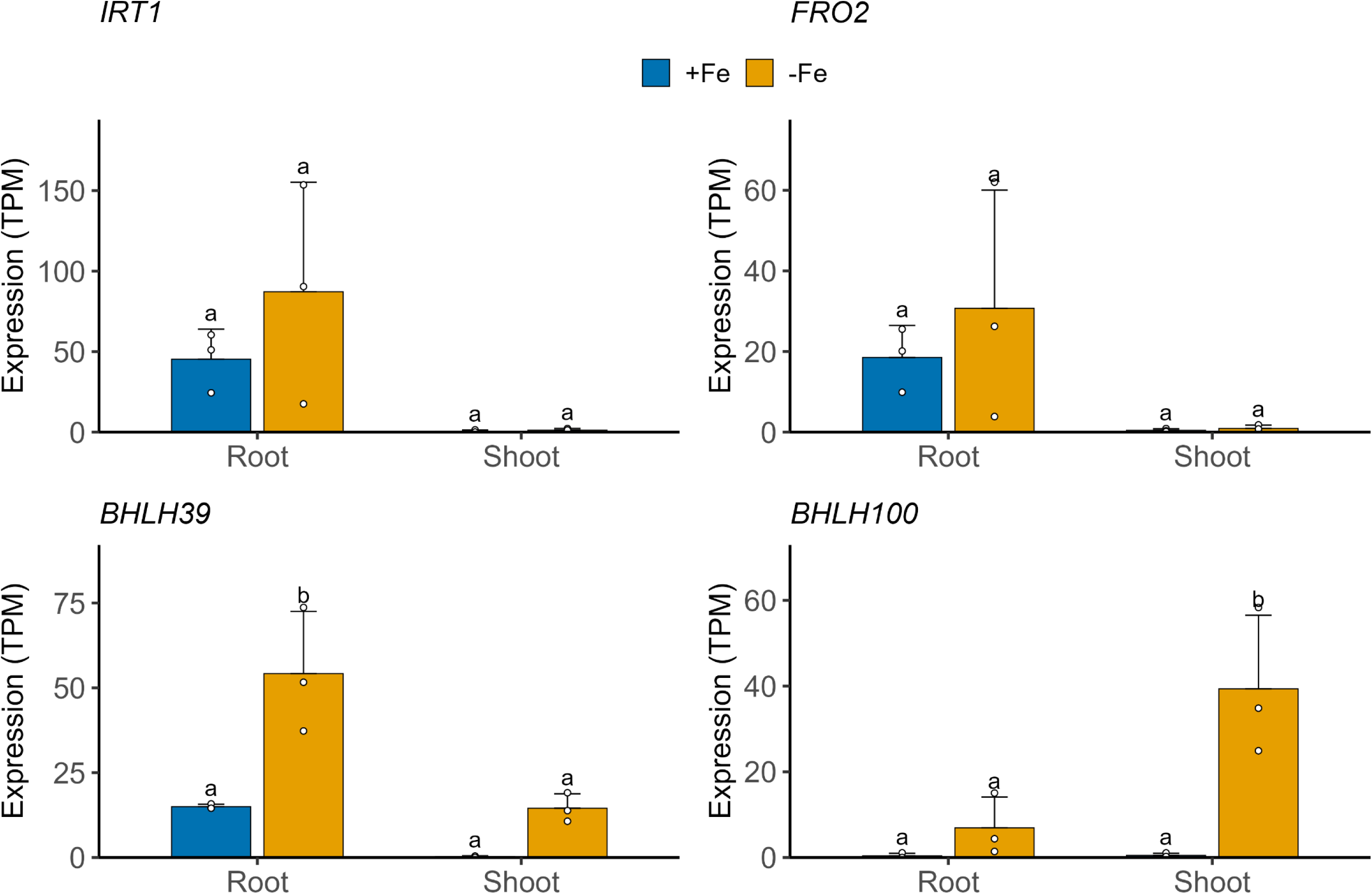
Validation of Fe deficiency treatment in wild-type plants. Expression of Fe deficiency marker genes in wild-type Col-0 roots and shoots under Fe-sufficient (+Fe) and Fe-deficient (−Fe) conditions. RNA-seq data are presented as transcripts per million (TPM). *IRT1* and *FRO2*, encoding the root Fe uptake transporter and ferric reductase, respectively, show higher expression under −Fe in roots, although the difference was not statistically significant due to replicate variability. *BHLH39* is induced under −Fe compared to +Fe in both roots and shoots. *BHLH100* is induced under −Fe compared to +Fe predominantly in shoots. These expression patterns confirm that the Fe deficiency treatment elicited the expected transcriptional −Fe response in wild-type plants. Data represent mean ± SD (n = 3 biological replicates). Statistical analysis was performed using one-way ANOVA followed by Tukey’s HSD test (p < 0.05). Different letters indicate statistically significant differences. Note that in the main figure heatmaps (Figure 3), these differences are not visible due to Z-score normalization being dominated by high expression values in *pb104::b104^dPAA* lines.

**Supplementary Figure S19.**
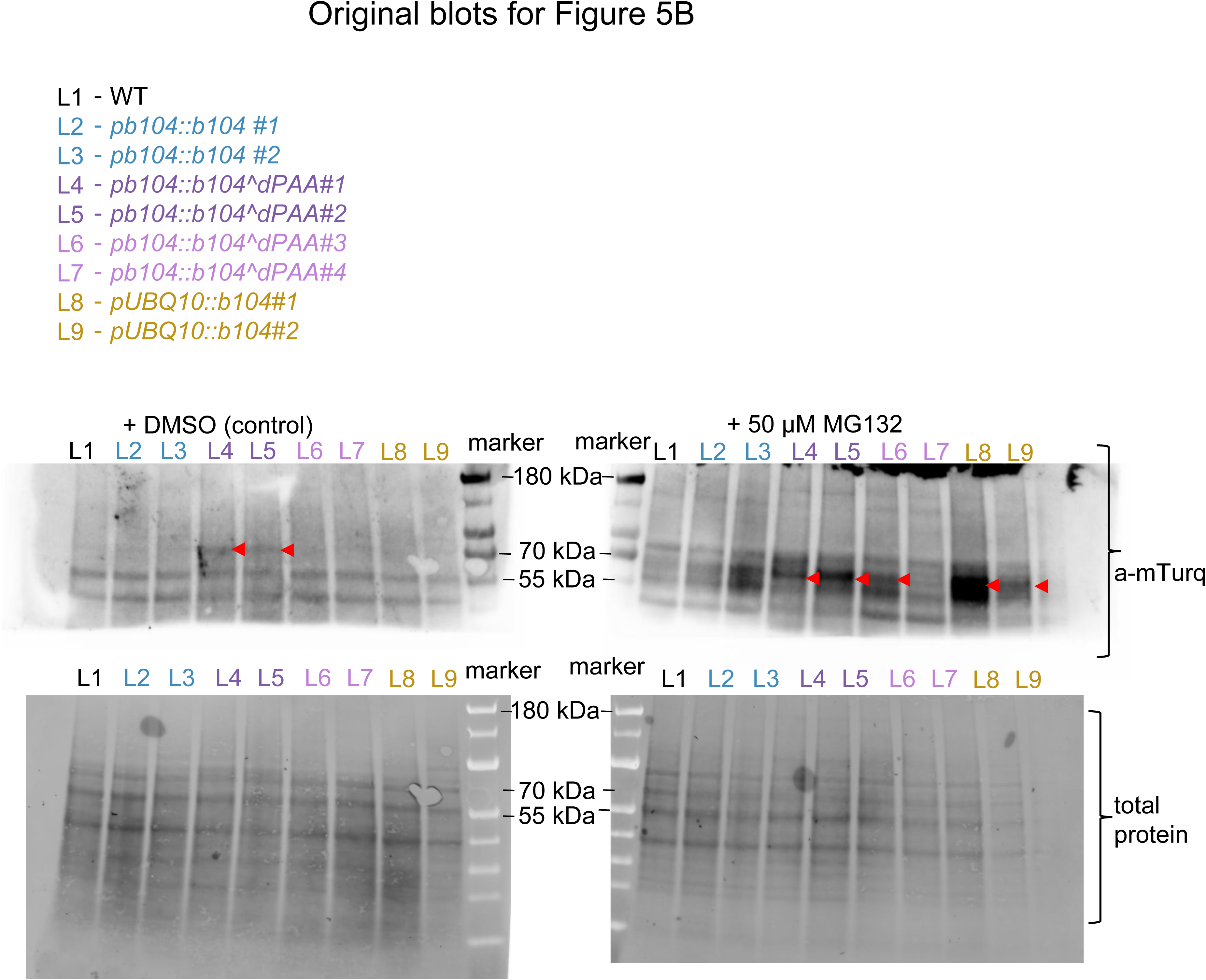
Original blots for Figure 5B.

**Supplementary Table 1.**
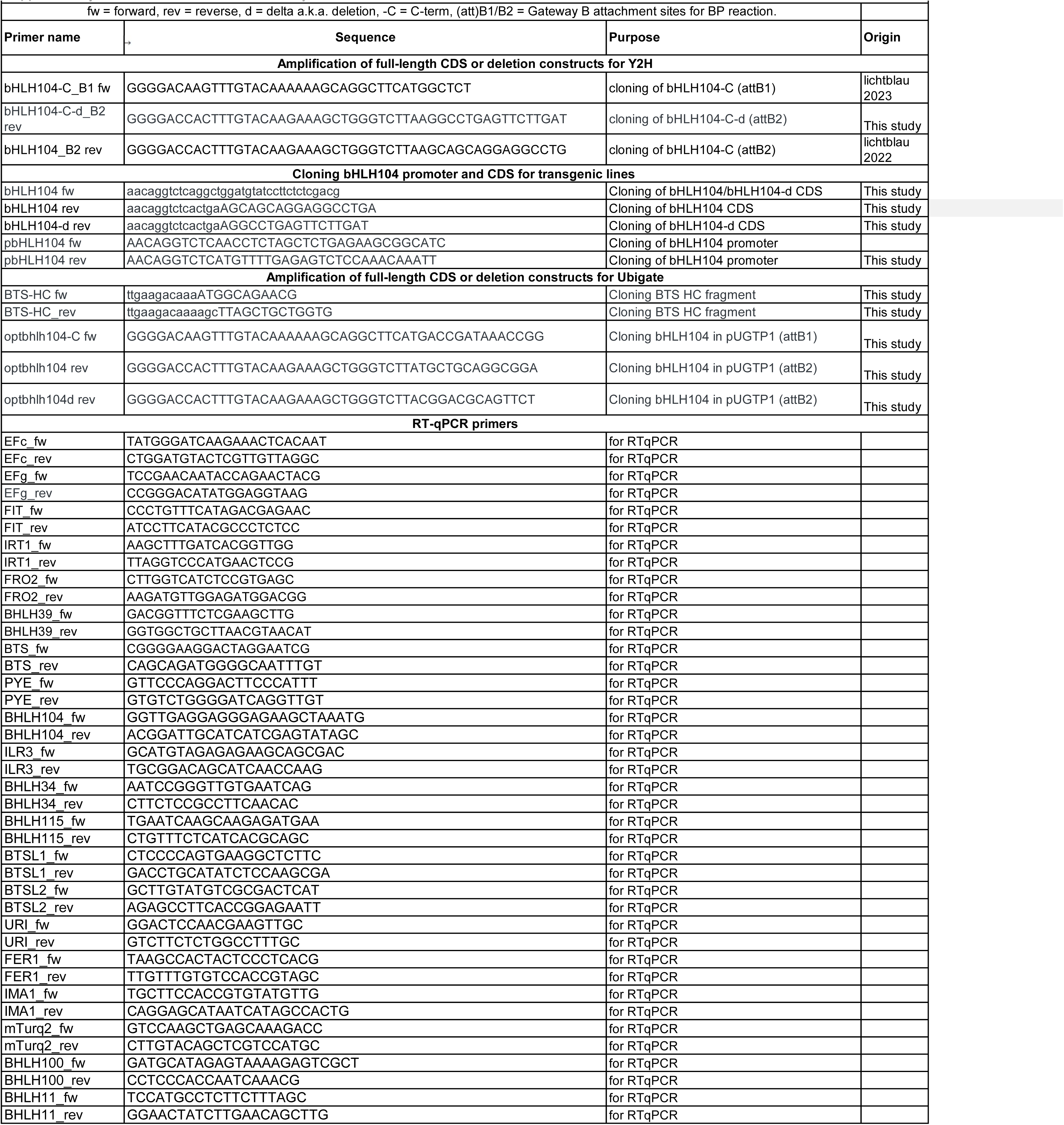
Primers used in this study.

